# Respiratory and C_4_-photosynthetic NAD-malic enzyme coexist in bundle sheath cells mitochondria and evolved via association of differentially adapted subunits

**DOI:** 10.1101/2021.06.16.448762

**Authors:** Meike Hüdig, Marcos A. Tronconi, Juan P. Zubimendi, Tammy L. Sage, Gereon Poschmann, David Bickel, Holger Gohlke, Veronica G. Maurino

## Abstract

In different lineages of Cleomaceae, NAD-malic enzyme (NAD-ME) was independently co-opted to participate in C_4_ photosynthesis. In the C_4_ Cleome species *Gynandropsis gynandra* and *Cleome angustifolia,* all NAD-ME genes (*NAD-MEα, NAD-MEβ1,* and *NAD-MEβ2*) were affected by C_4_ evolution and are expressed at higher levels than their orthologs in the C_3_ Cleome species *Tarenaya hassleriana*. In the latter C_3_ species, the NAD-ME housekeeping function is performed by two heteromers, NAD-MEα/β1 and NAD-MEα/β2, with similar biochemical properties. In both C_4_ species analyzed, this role is restricted the NAD-MEα/β2 heteromer. In the C_4_ species, NAD-MEα/β1 is exclusively present in the leaves, where it accounts for most of the enzymatic activity. GgNAD-MEα/β1 exhibits high catalytic efficiency and is differentially activated by the C_4_ intermediate aspartate, confirming its role as the C_4_-decarboxylase. During C_4_ evolution, GgNAD-MEβ1and CaNAD-MEβ1 lost their catalytic activity; their contribution to enzymatic activity results from a stabilizing effect on the associated α-subunit. We conclude that in bundle sheath cell mitochondria of C_4_ Cleome species, the functions of NAD-ME as C_4_ photosynthetic decarboxylase and as a tricarboxylic acid cycle-associated housekeeping enzyme coexist and are performed by isoforms that combine the same α subunit with differentially adapted β subunits.

## INTRODUCTION

Many of the world’s most productive crops are species bearing the C_4_ photosynthetic metabolism, as an adaptation to high light intensities, temperatures, and dryness (Sage et al., 2012). C_4_ plants have evolved biochemical pumps to concentrate CO_2_ at the site of ribulose-1,5-bisphosphate carboxylase/oxygenase (Rubisco), which decrease the oxygenation reaction and, thus, limit the wasteful flux through photorespiration (Furbank and Hatch, 1987). Compared with plants using the ancestral C_3_ photosynthetic pathway, C_4_ plants show higher water and nitrogen use efficiency, which allows increased productivity in warm habitats (Edwards et al., 2010). The transition from C_3_ to C_4_ photosynthesis involved complex alterations to leaf anatomy and biochemistry. This occurred in at least 66 linages of angiosperms independently and, thus, is an example of convergent evolution (Sage, 2016). Most C_4_ plants developed a spatial separation of the biochemical components of the CO_2_ pump by utilizing two adjacent cell types. In these plants, PEP-carboxylase is located in the cytosol of mesophyll cells, and the formed C_4_ intermediates are shuttled to the bundle sheath cells (BSCs) (Drincovich et al., 2011). The release of CO_2_ from malate in BSCs is mediated mainly by two different decarboxylases: NADP-malic enzyme (NADP-ME, EC 1.1.1.40), which is exclusively located to chloroplasts, and NAD-ME (EC 1.1.1.39), with exclusive location in the mitochondria (Maier et al., 2011; Wang et al., 2014). The majority of the C_4_ linages mainly use NADP-ME (Sage et al., 2011). The use of NAD-ME is mostly widespread in eudicot species, where it is found in approximately 20 C_4_ lineages (Sage et al., 2011).

The C_4_ enzymes did not evolve *de novo*. Instead, they were recruited from existing housekeeping isoforms (Aubry et al., 2011; Christin et al., 2013a; Maier et al., 2011; Monson, 2003). Most C_4_ enzymes originated by gene duplication of an existing non-photosynthetic isoform and the subsequent neofunctionalization of one gene copy (Alvarez et al., 2019; Christin et al., 2013b; Ludwig, 2016a; Saigo et al., 2004; Tausta et al., 2002). The C_4_-NAD-ME turned out to be an exceptional case. Recently, we described that the evolution of NAD-ME in higher plants is marked by sub-functionalization and differences in the frequency of gene duplication the two paralogous *α-* and *β-NAD-ME* gene lineages (Tronconi et al., 2020). Most angiosperm genomes maintained a 1:1 *α-NAD-ME*/*β-NAD-ME* relative gene dosage, but a significantly high proportion of species with C_4_-NAD-ME-type photosynthesis have a non-1:1 ratio of *α-NAD-ME*/*β-NAD-ME*. Specifically, some C_3_ and C_4_ species of Brassicales possess a single *NAD-MEα* gene and two *NAD-MEβ* genes (*β1* and *β2*) (Tronconi et al., 2020). In the independently evolved C_4_ species *Gynandropsis gynandra* and *Cleome angustifolia*, all three genes were affected by C_4_ evolution with the encoded NAD-MEβ1 subunits exhibiting several amino acids identically substituted and positively selected in both C_4_ species (Tronconi et al., 2020). Likely, these changes provided the basis for the recruitment of NAD-ME and its function in C_4_-physiology.

In its housekeeping role, plant NAD-ME is involved in mitochondrial malate respiration functioning as an associated enzyme to the tricarboxylic acid (TCA) cycle (Artus and Edwards, 1985; Fuchs et al., 2020; Grover et al., 1981; Tronconi et al., 2008). Plant NAD-ME mostly operates as a heteromer of α (∼ 63 kDa) and β (∼ 58 kDa) subunits that share about 65 % sequence identity (Tronconi et al., 2008), mirroring the existence of the paralogous *α-* and *β-NAD-ME* genes (Tronconi et al., 2020). Depending on the species and/or plant tissue, NAD-ME is active as a dimer, tetramer, and octamer (Artus and Edwards, 1985). In the C_3_ species *Arabidopsis thaliana*, NAD-ME is a heterodimer of the α- (*AtNAD-ME1,* At4g13560) and β-(*AtNAD-ME2*, At4g00570) subunits, while a small proportion of homodimers are present in components of the floral organ (Maier et al., 2011; Tronconi et al., 2018). The recombinant homodimers and the reconstituted heterodimer present similar catalytic efficiencies but differ in kinetic mechanisms and their regulation by metabolic effectors (Tronconi et al., 2008; Tronconi et al., 2010a; Tronconi et al., 2010b; Tronconi et al., 2015). It is intriguing that there are no references to housekeeping or photosynthetic isoforms of NAD-ME in any C_4_ plants. Due to the intricated molecular mechanism of evolution of NAD-ME (Tronconi et al., 2020), it is not surprising that despite its central importance in the C_4_-pathway, C_4_-NAD-ME has never been characterized at the molecular level. Even comparative transcriptomes of leaves of the closely related Cleome *Tarenaya hassleriana*, a C_3_ ornamental species, and *G. gynandra*, a NAD-ME C_4_ orphan crop species (Sogbohossou et al., 2018), did not reveal any C_4_-specific NAD-ME transcript (Brautigam et al., 2011).

Elucidation of the structural composition and biochemical characteristics of the components of the C_4_ pathway is not only important to gain knowledge on the evolutionary mechanisms underlying their origin but also because many efforts are currently under way to install characteristics of C_4_ photosynthesis in leaves of C_3_ crops (Ermakova et al., 2020; Kajala et al., 2011; Lin et al., 2020). This endeavor can only be fruitful, if it is built on a detailed understanding of the particular characteristics of the components of the C_4_ pathway. In this work, we addressed how NAD-ME in C_4_ plant mitochondria has been adapted to perform the housekeeping and C_4_ functions. We found that in C_4_ Cleome, the functions of NAD-ME as a respiratory enzyme and specific photosynthetic decarboxylase are performed by separated enzymatic entities originating from the combination of the α-subunit with differentially evolved β-subunits.

## Results

### Expression of *NAD-ME* genes in C_3_ and C_4_ Cleome species

To find candidate gene(s) of C_4_-NAD-ME in Cleome, we analyzed the diel expression pattern of all *NAD-ME* genes (*α*, *β1,* and *β2*) in fully expanded leaves of *T. hassleriana* (C_3_) and *G. gynandra* (C_4_) by quantitative transcriptional analysis (Figure 1 and Supplemental Table 1).

**Figure 1.**
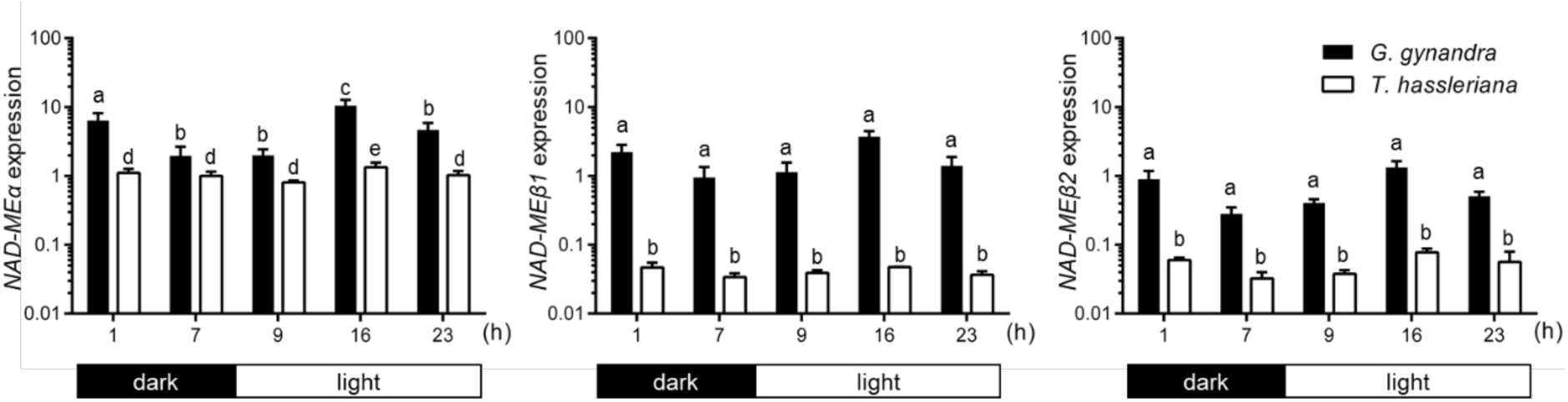
Expression levels of genes encoding NAD-ME proteins in *G. gynandra* and *T. hassleriana* relative to *ACT2* in leaves of 5-week-old plants over a full day. The values represent the mean ± SE; n=three independent replicates. Within species, same letters indicate no statistical differences, different letters indicate statistical differences among the time points based on two-way ANOVA (α=0.05) with a post-hoc Tukey test (see results of multiple comparisons with confidence intervals and significance levels in Supplemental Table 1).

The expression patterns of the *NAD-MEβ* genes do not change drastically during a diurnal cycle in the C_3_ and C_4_ species (Figure 1). In both species, the expression of the *NAD-MEα* genes slightly peaks in the middle of the light period (Figure 1). We found that *GgNAD-ME*α is expressed 2- to 10-fold higher than *GgACT2*, while *GgNAD-MEβ1* reaches up to 3.7-fold and *GgNAD-MEβ2* up to 1.3-fold higher expression than *GgACT2* at midday (Figure 1). *ThNAD-ME*α expression levels are similar to that of *ThACT2*, while *ThNAD-MEβ1* and *ThNAD-MEβ2* show approx. 20-fold lower expression than the control gene (Figure 1). As all three *GgNAD-ME* genes are expressed at much higher levels than their homologs in *T. hassleriana*, we cannot unambiguously identify a concrete *C_4_-NAD-ME* candidate gene by the transcriptional analysis.

### NAD-ME entities in photosynthetic and heterotrophic organs of C_3_ and C_4_ Cleome

We explore the presence of NAD-ME isoforms in photosynthetic and heterotrophic organs of C_3_ (*T. hassleriana*) and C_4_ (*G. gynandra* and *C. angustifolia*) Cleome species by coupling *in gel* NAD-ME activity assay after gradient native PAGE with the identification of NAD-ME subunits in the gel slices through mass spectroscopy (MS). Overall, *in gel* activity assays indicated different protein bands showing NAD-ME activity distributed over species and different organs (migration levels 1 to 4 in Figure 2).

**Figure 2.**
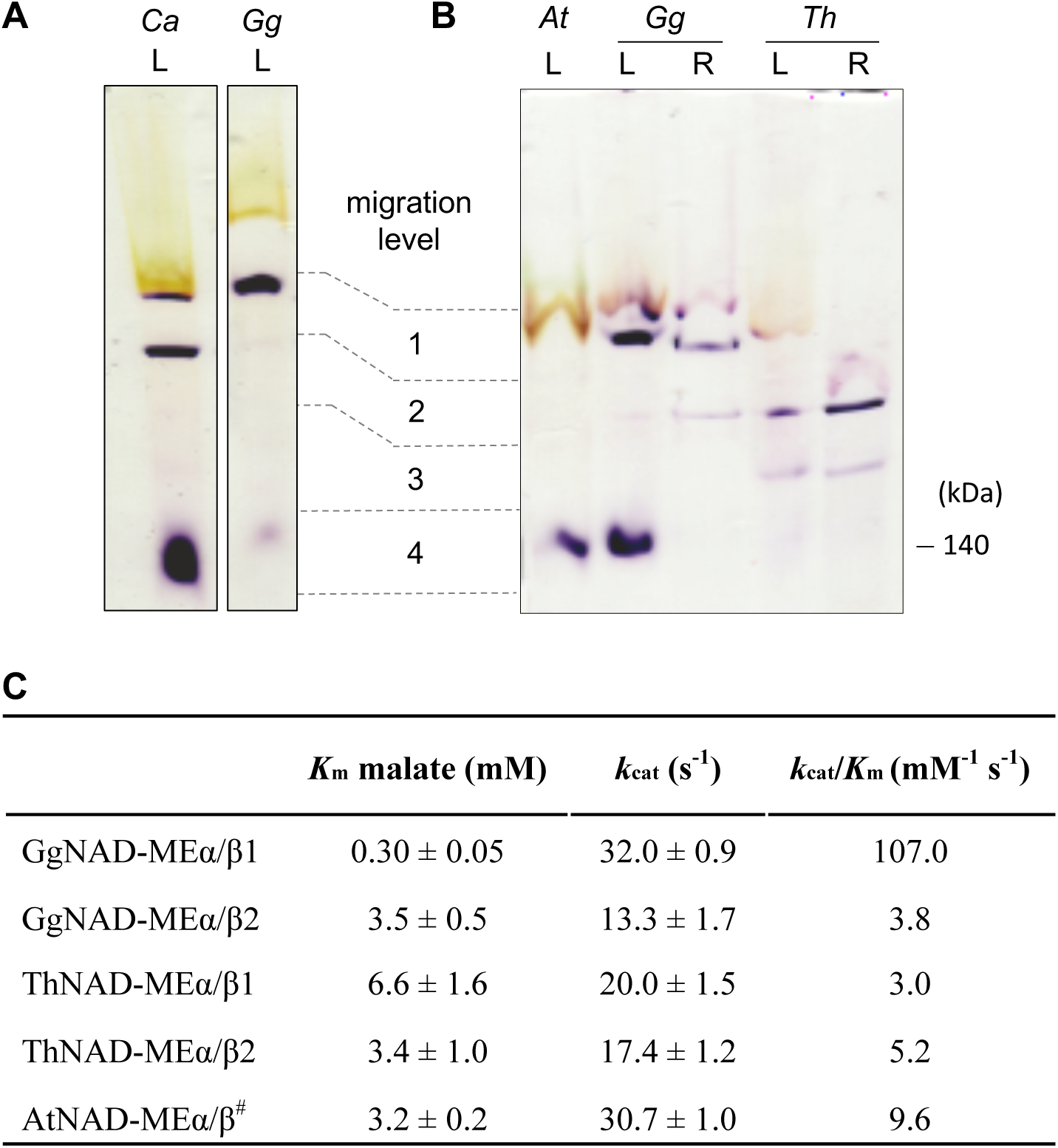
Behavior of NAD-ME entities of Cleome species. (A) Gradient native PAGE coupled to NAD-ME activity assay of soluble protein extracts of leaves (L, 40 µg protein) of *C. angustifolia* (Ca) and *G. gynandra* (Gg). (B) Gradient native PAGE coupled to NAD-ME activity assay of soluble protein extracts of leaves (L, 40 µg protein) and roots (R, 20 µg protein) of *G. gynandra* (Gg), and *T. hassleriana* (Th); *A. thaliana* (At) soluble protein extracts of leaves ere used as control (40 µg protein) (Tronconi et al., 2008). For the *in gel* activity assays in A and B, the gels were incubated during 150 min in the assay medium at pH 6.8. A violet precipitate indicates NAD-ME activity. Migration levels 1 to 4 are indicated on the right. The molecular weight of the control AtNAD-ME (Tronconi et al., 2008) is indicated on the left. (C) Kinetic parameters of recombinant NAD-ME entities identified in protein extracts of *G. gynandra* and *T. hassleriana*. Kinetic data were fitted by nonlinear regression. ^#^Data taken from (Tronconi et al., 2010b). Values represent mean ± SE; n=three independent enzyme preparations, each measured in triplicate. As GgNAD-MEβ1 is not active, *k*_cat_ was calculated assuming the formation of a heterodimer with only one active site in GgNAD-MEα/β1.

Leaves of *G. gynandra* and *C. angustifolia* show two main protein bands with NAD-ME activity of very different mobilities (migration level 1 and 4, Figure 2A); these bands are not present in the C_3_ species (Figure 2B). One protein band with NAD-ME activity has a very high mobility (migration level 4, Figure 2A) and showed the presence of only NAD-MEα and NAD-MEβ1 after MS analysis (Supplemental Table 2). As the mobility of this protein band is similar to that of the heterodimer NAD-MEα/β found in *A. thaliana* leaves (Tronconi et al., 2008) (Figure 2B), we propose that it corresponds to a NAD-MEα/β1 heterodimer. The second protein band found exclusively in leaves of C_4_ species has very low mobility (migration level 1, Figure 2A) and appears rapidly after incubation in the activity assay buffer (Supplemental Figure 1A). MS analysis of this low mobility protein band from both C_4_ species showed the presence of all three NAD-ME subunits (Supplemental Tables 2). Most likely, two different heteromers with a similar velocity of migration, NAD-MEα/ß1 and NAD-MEα/ß2, contribute to the NAD-ME activity of this low mobility band of leaves of the C_4_ species.

We further analyzed roots of the C_4_ species *G. gynandra*, which show a predominant NAD-ME activity band with low mobility at migration level 1 (Figure 2B). MS analysis of this protein band identified peptides corresponding to NAD-MEα and NAD-MEβ2 (Supplemental Table 2), indicating that the NAD-MEα/β2 heteromer is the only isoform present in roots.

Leaves and roots of the C_3_ plant *T. hassleriana* show a protein band with NAD-ME activity that is not found in organs of the C_4_ species (migration level 3, Figure 2B). This protein band has a slightly lower mobility than that of the *A. thaliana* NAD-ME heterodimer. MS analysis of the protein bands indicated the presence of NAD-MEα and identified peptides that correspond to both NAD-ME β-subunits (Supplemental Table 2). Thus, we suggest that the NAD-MEα/β1 and NAD-MEα/β2 isoforms are present in the protein bands of leaves and roots and have very similar electrophoretic mobilities.

The three analyzed species, G. *gynandra*, *C. angustifolia*, and *T. hassleriana,* share an additional protein band with NAD-ME activity and intermediate mobility (migration level 2, Figure 2A and B). MS analysis indicated the presence of a NADP-ME in all these protein bands, which was confirmed by *in gel* NADP-ME activity assay after native PAGE (Supplemental Figure 1B). As the identified NADP-MEs (Th09755, Th20644 and Gg17666) are homologues of co-factor promiscuous cytosolic AtNADP-MEs (Gerrard Wheeler et al., 2008), we conclude that the band at migration level 2 found in most samples represents a NADP-ME that also uses NAD as a cofactor at least *in vitro*.

Taken together, leaves and roots of the Cleome C_3_ species *T. hassleriana* most likely possess two NAD-ME isoforms, ThNAD-MEα/β1 and ThNAD-MEα/β2, formed by the alternative combination of the α-subunit with either of the β-subunits. The Cleome C_4_ species *G. gynandra* and *C. angustifolia* also possess two NAD-ME isoforms. Ca/GgNAD-MEα/β2 is present in heterotrophic and photosynthetic organs and is assembled as a heteromer of low mobility. The second isoform is exclusively present in photosynthetic tissues and is formed by the association of the α- and β1-subunits. In the conditions of our assay, Ca/GgNAD-MEα/β1 is responsible for the major part of NAD-ME activity and is active in two structural assemblies: as heterodimer and as a higher-order heteromer.

### Biochemical properties of recombinant NAD-ME of C_3_ and C_4_ Cleome

To analyze if the identified NAD-ME entities in C_4_ Cleome differ in biochemical properties, we performed *in vitro* analyses with GgNAD-MEs. We co-expressed the mature GgNAD-MEs subunits in *E. coli* and purified them using the His-Tag fused to only one subunit (Supplemental Figure 2). That way, the co-elution of the subunits indicates specific interactions of the proteins. We found that GgNAD-MEα interacts with the NAD-MEβ subunits to form the GgNAD-MEα/β1 and GgNAD-MEα/β2 heteromers, while the recombinant GgNAD-MEβ1 and GgNAD-MEβ2 subunits do not interact with each other (Supplemental Figure 2). These results reinforced the results of *in gel* NAD-ME assay coupled to MS analysis, showing that in leaves of C_4_ Cleome NAD-ME forms two heteromers constituted by the association of an α-subunit with a different β-subunit (GgNAD-MEα/β1 and GgNAD-MEα/β2, Figure 2A).

The recombinant GgNAD-MEα/β1 and GgNAD-MEα/β2 are active enzymes with a similar optimum pH of 6.2-6.4 and K_m,NAD_ of 0.6-0.7 mM, but different kinetic parameters with respect to the substrate malate. GgNAD-MEα/β2 has kinetic parameters in the same range as that of the Arabidopsis isoform (Figure 2C). In contrast, GgNAD-MEα/β1 has an approx. 30 times higher catalytic efficiency than GgNAD-MEα/β2 (Figure 2C). This high catalytic efficiency results from a high apparent affinity to malate, which is 12 times greater than that of GgNAD-MEα/β2, and a turnover rate 2-times higher than that of GgNAD-MEα/β2 (Figure 2C). Similarly, we also expressed and analyzed the NAD-ME isoforms identified in the C_3_ species *T. hassleriana* (ThNAD-MEα/β1 and ThNAD-MEα/β2). We found that both isoforms have similar kinetic parameters to those of GgNAD-MEα/β2 and AtNAD-MEα/β (Figure 2C), indicating their housekeeping functions in mitochondrial respiration.

In gradient native PAGE, the recombinant GgNAD-MEα/β1 is found as two bands of different electrophoretic mobilities, which let us assume that in the conditions of the assay, this isoform can be found as a heterodimer and heterotetramer (Supplemental Figure 3), as also observed with extracts of Cleome organs (Figure 2A). Consistently, GgNAD-MEα/β1 eluted into two peaks after gel filtration chromatography: a peak representing the dimer (molecular mass of 132 ± 15 kDa) and a second broad peak representing proteins of high molecular weight (between 250 and 460 kDa). Recombinant GgNAD-MEα/β2 eluted as a broad peak consistent with a high molecular weight protein assembly (between 250 and 460 kDa).

Taken together, the analysis of recombinant NAD-ME proteins indicates that in *G. gynandra*, the association of the α- and β1-subunits renders a NAD-ME isoform, GgNAD-MEα/β1, with high affinity for malate and high catalytic efficiency. These are expected properties for an enzyme to fulfill a role in a high-flux metabolic pathway, such as the C_4_ photosynthetic pathway. On the other hand, the housekeeping NAD-ME of *G. gynandra* is formed by the association of α- and β2-subunits, GgNAD-MEα/β2. The C_3_ species *T. hassleriana* possesses two housekeeping NAD-ME isoforms with similar biochemical properties, which are formed by the alternative combination of the α-subunit with either of the β-subunits, ThNAD-MEα/β1 and ThNAD-MEα/β2.

### NAD-MEβ1 evolved as a non-catalytic subunit in C_4_ Cleome

To aid in understanding the molecular determinants of the evolution of different NAD-ME entities in Cleome, we analyzed the single NAD-ME proteins of *G. gynandra* and *T. hassleriana* produced *in vitro* (Figure 3A; Supplemental Figure 4). The results indicated that all recombinant proteins, except for one, are active enzymes (Figure 3A). Intriguingly, we found that GgNAD-MEβ1 is the only inactive protein. Similarly, we found that the recombinant NAD-MEβ1 of *C. angustifolia* is also inactive.

**Figure 3.**
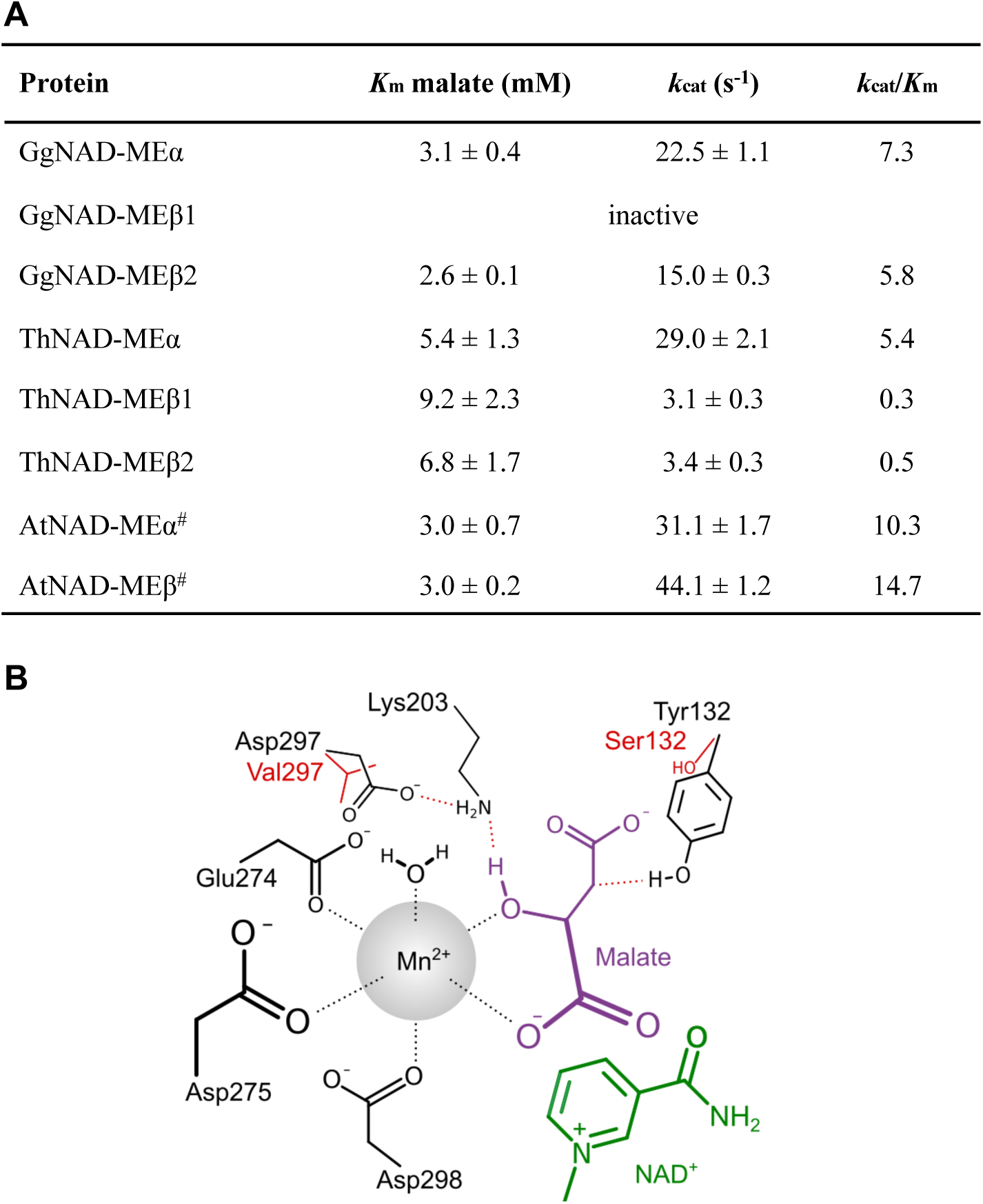
NAD-MEβ1 evolved as a non-catalytic subunit in C_4_ Cleome. (A) Kinetic parameters of recombinant NAD-ME single proteins from *G. gynandra, C. angustifolia,* and *T. hassleriana*. Kinetic data were fitted by nonlinear regression. Values represent mean ± SE of at least three independent enzyme preparations. ^#^Data taken from (Tronconi et al., 2008). (B) Schematic representation of the active site of GgNAD-MEβ1. The substitutions in GgNAD-MEβ1 with respect to GgNAD-MEβ2 are highlighted in red.

Recently, we reported that NAD-MEβ1 from the C_4_ species *G. gynandra* and *C. angustifolia* possess five identically substituted amino acids (V131, S132, H195, V297, and K605) compared with other species and the β2-NAD-MEs of the same species (I131, Y132, Q195, D297, and E605, sequence numbering after alignment to the β1 sequence) (Tronconi et al., 2020). To analyze the structural implications of the substitutions in C_4_-NAD-MEβ1, we built homology models of the α, β_1_, and β_2_-isoforms of *G. gynandra* and *A. thaliana*. Among the five conserved substitutions in the C_4_-NAD-MEβ1 isoforms, only two lead to major changes in the side chain properties: Y132S and D297V. Both substitutions are located near the catalytic center of the enzyme (Figure 3B). Y132 is directly involved in the catalytic mechanism of the enzyme, donating a proton to the enolpyruvate intermediate to facilitate tautomerization to pyruvate (Chang and Tong, 2003). For that, the hydroxy group needs to be close to the C3 atom of malate. We found that the Y132S substitution almost doubles the distance between the malate C3 and the hydroxy group (3.7 Å to 6.8 Å) (Figure 3B). Furthermore, the pK_a_ of serine is ∼ 3 log units larger than that of tyrosine. Thus, serine at position 132 likely cannot donate a proton to enolpyruvate. D297 is not directly involved in the catalytic mechanism but forms a charge-assisted hydrogen bond with K203; the latter is not protonated and accepts the proton from the 2-OH group of malate in the oxidation step (Chang and Tong, 2003). Due to the loss of the polar interaction through the D297V substitution, likely, K203 becomes more mobile, and its pK_a_ decreases, which weakens its basicity. Hence, we propose that the substitutions Y132S and D297V likely affect the (de-)protonation events that occur during the catalytic cycle of the NAD-ME. This supports that NAD-MEβ1 evolved as a non-catalytic subunit in the C_4_ Cleome species *G. gynandra* and *C. angustifolia*.

### GgNAD-MEβ1 confers regulatory functions to the photosynthetic NAD-ME

The evolutionary changes that occurred in the sequence of NAD-MEβ1 in C_4_ Cleome species (Tronconi et al., 2020) let us hypothesize that this non-catalytic subunit was retained because it must have evolved other molecular properties important for its function in C_4_ photosynthesis. One such function could be the regulation of the enzymatic activity by metabolic effectors.

To test this hypothesis, we analyzed the influence of selected metabolites on the activity of GgNAD-ME isoforms. From the glycolytic and TCA cycle intermediates analysed, we found that GgNAD-MEα/β1 activity is strongly enhanced by fructose-1,6-bisphosphate (FBP), phosphoenolpyruvate (PEP), fumarate, and acetyl-CoA (Figure 4A). This agrees with the reported regulation of all *A. thaliana* NAD-MEs (Tronconi et al., 2010a). Also, GgNAD-MEα is strongly activated by fumarate, and GgNAD-MEβ2 is activated by FBP, PEP, and CoA (Figure 4A). As we already showed that FBP, PEP, and CoA bind to the surface of Arabidopsis NAD-MEβ subunit (Tronconi et al., 2010a), the activation of GgNAD-MEα/β1 by these effectors provides evidence that GgNAD-MEβ1, although non-catalytic *in vitro*, is likely a subunit with conserved regulatory functions. Furthermore, GgNAD-MEα/β1 shows specific inhibition by 3PG and activation by OAA (Figure 4A).

**Figure 4.**
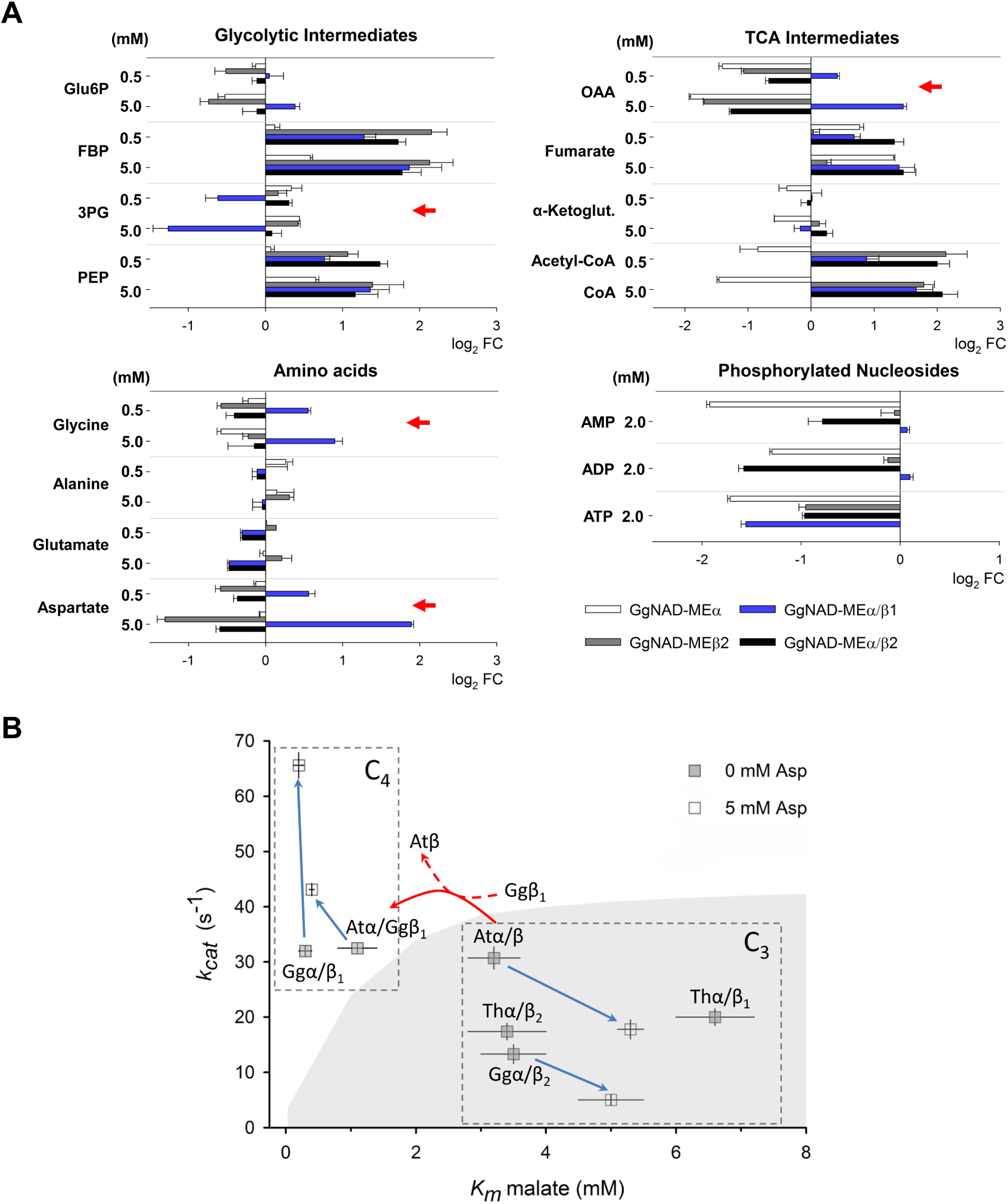
Regulatory properties of recombinant GgNAD-MEs. (A) Modulation of the activity of GgNAD-ME entities by selected metabolites. Enzymatic activities were measured in the absence or presence of each effector. Malate concentration was kept at a value of *K*_m_/3 for each enzyme. The results are presented as the log_2_ FC (FC, fold change = enzymatic activity in the presence of each effectors/enzymatic activity measured in the absence of the effectors). FC values lower than 0.7 (log_2_ < −0.5) or higher than 1.3 (log_2_ > 0.4) were significantly different to 1 (p-value < 0.05, One Way ANOVA with Holm-Sidak method) and considered as an inhibition (left bars) or activation (right bars) of the enzymatic activity, respectively. Red arrows indicate metabolites that differentially regulated GgNAD-ME α/β1 activity. (B) Cartesian plane representation of the kinetic parameters of recombinant GgNAD-MEα/β1 (Ggα/β1), GgNAD-MEα/β2 (Ggα/β2), ThNAD-MEα/β1 (Thα/β1), ThNAD-MEα/β1 (Thα/β2), AtNAD-MEα/β (Atα/β) and the interspecies chimera AtNAD-MEα/GgNAD-MEβ1 (Atα/Ggβ1) in the absence and the presence of aspartate. Mean values of *K*_m_ malate and *k*_cat_ are shown as an ordered pair (*K*_m_, *k*_cat_) with error bars indicatig ± SE. The blue arrows connect the kinetic parameters values in the absence of aspartate (gray squares) and those obtained in the presence of 5 mM aspartate (white squares). The red arrow indicates the replacement of the β-subunit which is associated with the Atα-subunit in the interspecies chimera Atα/Ggβ1. The gray area separate enzymes with *k*_cat_/*K*_m_ values lower than 100 s^−1^*mM^−1^. All points on the boundary line have a constant *k*_cat_/*K*_m_ value (= 100). The boundary line was generated from the M-M equation in the form rate=(k_cat_/K_m_)E_0_[S]/(1+[S]/K_m_) at E_0_ = 1 (arbitrary units) and [S] = 0.5 mM (S: malate), setting rate as dependent and *K*_m_ as independent variable, respectively, varying *K*m from 0.02 to 8 mM.

From the amino acid tested, glycine and aspartate specifically stimulated GgNAD-MEα/β1 activity with apparent constants of activation (A_50_) of 6.2 mM for glycine and 3.8 mM for aspartate. Aspartate has a stronger inhibitory effect on the activity of GgNAD-MEβ2 and GgNAD-MEα/β2 than glycine and does not modify GgNAD-MEα activity (Figure). Interestingly, in GgNAD-MEβ2, aspartate behaved as a competitive inhibitor of malate (Supplemental Table 3), suggesting that it binds to the active site of the β-subunit.

Given that glycine and aspartate were not previously described as regulators of any plant NAD-ME, we comparatively analyzed their effects on the activities of GgNAD-MEα/β1, AtNAD-MEα/β, and the interspecies chimera AtNAD-MEα/GgNAD-MEβ1. Glycine had no effect on the activities of AtNAD-MEα/β and AtNAD-MEα/GgNAD-MEβ1. However, aspartate also activates AtNAD-MEα/GgNAD-MEβ1 (Figure 4B and Supplemental Table 3) with an A_50_ value of 4.5 mM. In the presence of 5 mM aspartate, the catalytic efficiencies of GgNAD-MEα/β1 and AtNAD-MEα/GgNAD-MEβ1 increase up to 3-fold due to increases in both malate affinity and turnover rate (Figure 4B and Supplemental Table 3). Even more, in the absence of aspartate, the affinity for malate of AtNAD-MEα/GgNAD-MEβ1 is 3-fold higher than that of AtNAD-MEα/β (Figure 4B and Supplemental Table 3). Aspartate decreases the enzymatic efficiency of AtNAD-MEα/β by a similar magnitude as in GgNAD-MEβ2 and GgNAD-MEα/β2 (Figure 4B and Supplemental Table 3).

All these results indicate that (i) the GgNAD-MEβ1 subunit is involved in the specific kinetic properties of GgNAD-MEα/β1, high affinity for malate and turnover rate, and activation by aspartate; (ii) in the β-subunits aspartate most probably binds to the active site and exerts different regulatory effects depending on the evolutionary adaptations of the subunits; (iii) despite the evolutionary divergence between species, the interaction of the NAD-MEα- and β-subunits is largely conserved; and (iv) the association of the non-catalytic GgNAD-MEβ1 subunit to the paralogous α-partner of Arabidopsis renders a NAD-ME entity with similar properties as GgNAD-MEα/β1.

### GgNAD-MEβ1 stabilizes an associated NAD-MEα subunit

To scrutinize how the non-catalytic GgNAD-MEβ1 subunit influences the activity of an associated GgNAD-MEα subunit, we performed molecular dynamics simulations of four dimeric NAD-ME with aggregate simulation times of 2.5 μs each (Supplemental Figure 5). We combined a “catalytic” NAD-MEα subunit with “effector” subunits α, β1, and β2, resulting in the dimers GgNAD-MEα/α, GgNAD-MEα/β1, GgNAD-MEα/β2, and ThNAD-MEα/β1. Then, we subjected the conformational ensembles to a Constraint Network Analysis to reveal the hierarchy of structural rigidity of the dimers. We focused on the effects of the “effector” subunits on the “catalytic” NAD-MEα subunit.

A comparison of the chemical potential energies due to non-covalent bonding per residue (see Eq. 1 Material and Methods) for the “catalytic” NAD-MEα subunits showed that the GgNAD-MEβ1 subunit has a more stabilizing influence on the associated GgNAD-MEα subunit than any of the other “effector” subunits (Figure 5A and B, Supplemental Figure 6). We found that the more stabilizing influence of GgNAD-MEβ1 originates from forming more rigid contacts across the interface to the GgNAD-MEα subunit than any of the other “effector” subunits (Figure 5C, Supplemental Figure 6B). This stabilizing effect percolates from the interface region to the active site (Figure 5B and C, Supplemental Figure 6A). The higher structural stability of the active site likely improves the binding of the substrate and cofactor or their orientation, therefore, leading to the higher catalytic efficiency of GgNAD-MEα/β1 with respect to all other NAD-ME entities analyzed (Figure 2B). The same mechanism may lead to the increased catalytic efficiency of the interspecies chimera AtNAD-MEα/GgNAD-MEβ1 with respect to the AtNAD-MEα/β dimer (Figure 4B and Supplemental Table 3).

**Figure 5.**
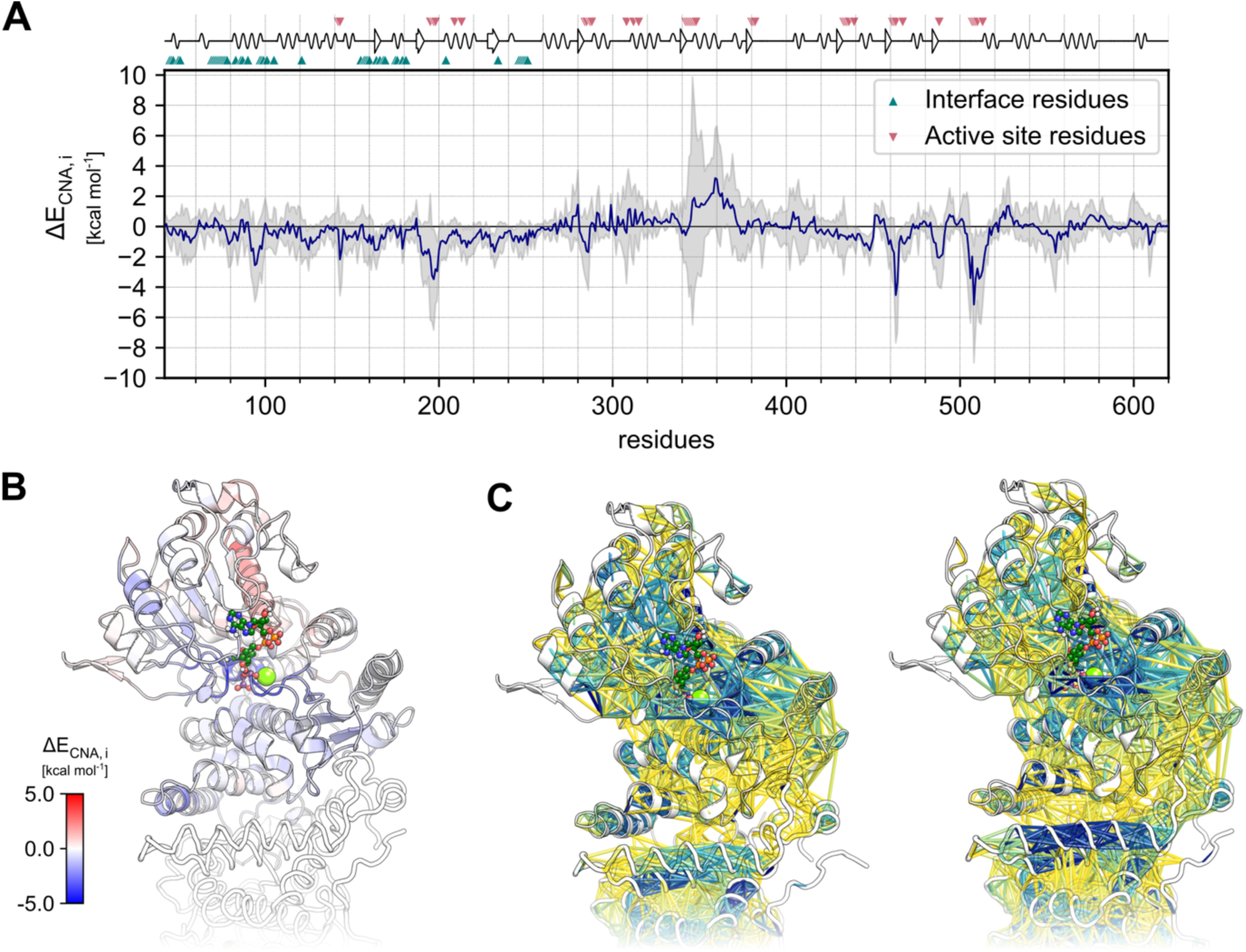
(A) The difference profile of the chemical potential energy due to non-covalent bonding per residue of the GgNAD-MEα subunits of a GgNAD-MEα/α homodimer and a GgNAD-MEα/β1 heterodimer. Values < 0 indicate a stabilizing effect of the β1 subunit on the α subunit with respect to the homodimer, values > 0 a destabilizing effect. Above the plot, a schematic representation of the enzymes’ secondary structure is shown, with markers below highlighting residues in the subunit-interface and markers above highlighting residues in the active site. (B) The difference profile from A mapped onto the “catalytic” α subunit of an GgNAD-MEα/β1 heterodimer (with bound NAD^+^ and L-malate) with the “effector” subunit being depicted as a white ribbon. Blue (red) colors indicate more (less) structurally stable regions than in the GgNAD-MEα/α homodimer. Corresponding representations of all analyzed systems are shown in Supplemental Figure 6A. (C) Depiction of rigid contact strength between amino acid pairs in the GgNAD-MEα/α homodimer (left) and the GgNAD-MEα/β1 heterodimer (right) with bound NAD^+^ and L-malate. A rigid contact denotes that two residues are part of a structurally rigid cluster in the structure. Blue (yellow) colors indicate more (less) prevalent rigid contacts. The “catalytic” α subunit is depicted as helices and sheets at the top, the “effector” subunit as ribbons below. In the GgNAD-MEα/β1 heterodimer, there are more rigid contacts across the interface than in the GgNAD-MEα/α homodimer.

Furthermore, we investigated how specific substitutions in the NAD-MEβ1 subunit in C_4_ Cleome may stabilize an associated NAD-MEα subunit. For this, we applied an ensemble-based perturbation approach implemented in constrained network analysis, perturbing selected residues in GgNAD-MEβ1 *versus* the corresponding residues in GgNAD-MEβ2. We applied the approach on four substitution sites that we previously showed to be identically substituted in *C. angustifolia* and *G. gynandra* NAD-MEβ1 with respect to other NAD-MEβ enzymes: I131V, Y132S, Q195H, and D297V (Tronconi et al., 2020). We found that, although the I131V substitution shows no impact on the rigidity of GgNAD-MEβ1, all other substitutions reduce the stability around the inactive catalytic site (Supplemental Figure 7). This destabilization, however, remains localized in the NAD-MEβ1 subunit. Interestingly, the Q195H substitution additionally exhibits a stabilizing effect in the interface region, which also percolates to the bound NAD-MEα subunit (Supplemental Figure 7C). These results suggest that H195 may have evolved in *C. angustifolia* and *G. gynandra* NAD-MEβ1 to stabilize the associated subunit.

### Immunolocalization studies

Our results indicate that in C_4_ Cleome, a housekeeping NAD-MEα/β2 and photosynthetic NAD-MEα/β1 (C_4_-NAD-ME) isoforms coexist in photosynthetic tissues and are formed by a differential combination of subunits. This gives rise to two hypotheses regarding the cellular location of both enzymatic entities in photosynthetic tissues of C_4_ Cleome. Either housekeeping and C_4_-NAD-ME are separated by cell type, or the housekeeping NAD-ME is found in every cell type, and only C_4_-NAD-ME is exclusively located in BSCs. To answer this question, we conducted immunolocalization analysis using leaves of *G. gynandra* and specific antibodies raised against each of the three NAD-ME subunits. We found high density labelling of GgNAD-MEα and GgNAD-MEβ2 subunits in mitochondria of MSC (Figure 6). The use of antibodies against GgNAD-MEβ1 rendered few spots, indicating either very low occurrence of the NAD-MEβ1 subunit in MC mitochondria or just background labelling. Mitochondria of BSC presented immunolabelling of all three subunits and at much higher density than those in MC (Figure 6 and Supplemental Figure 8). These results indicate that the housekeeping and photosynthetic NAD-ME entities coexist in BSC mitochondria of the C_4_ Cleome species (Figure 7).

**Figure. 6.**
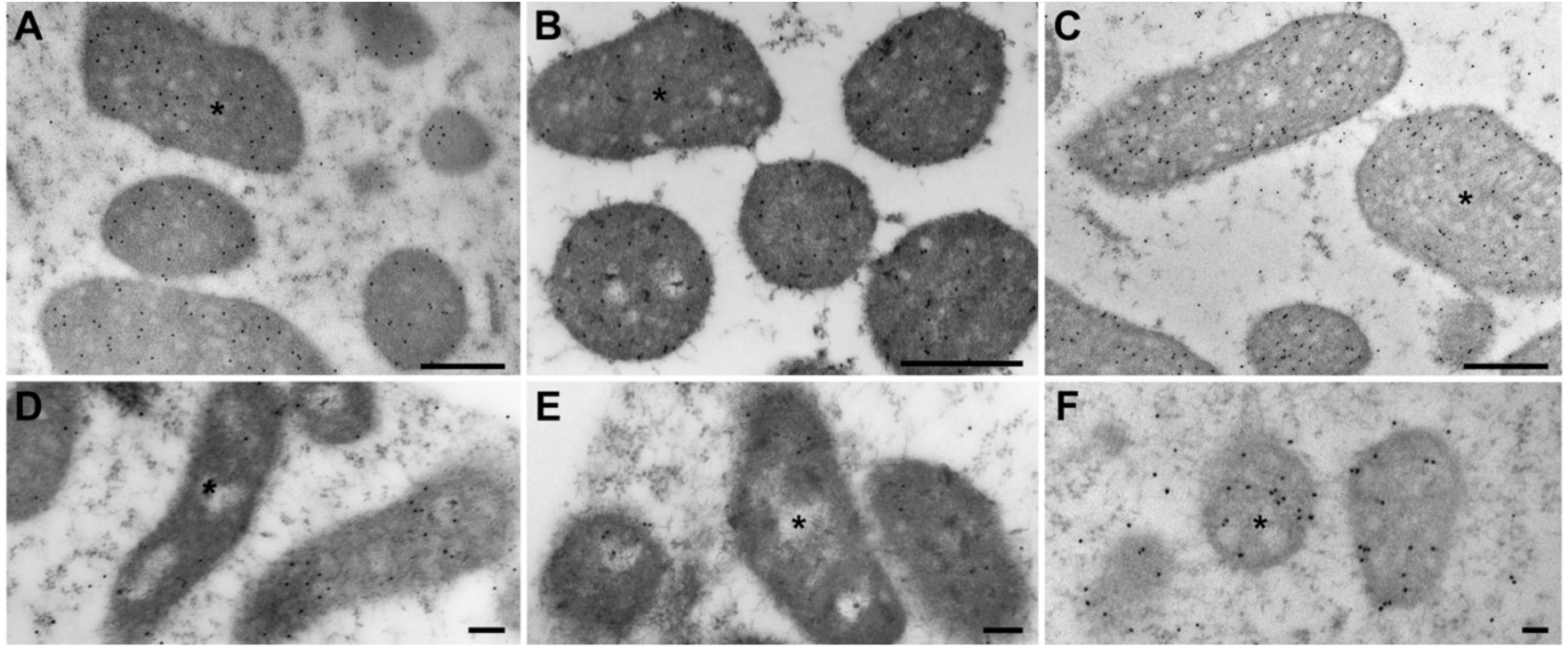
Transmission electron micrographs of bundle sheath (A-C) and mesophyll (D-F) cells of *Gynandropsis gynandra* showing mitochondria with immunogold labelling (black particles) of GgNAD-MEα (A, D), GgNADMEβ1 (B, E), and GgNAD-MEβ2 (C, F). *, mitochondrion. Bars, 500 nm (A-C); 100 nm (D-F).

**Figure 7.**
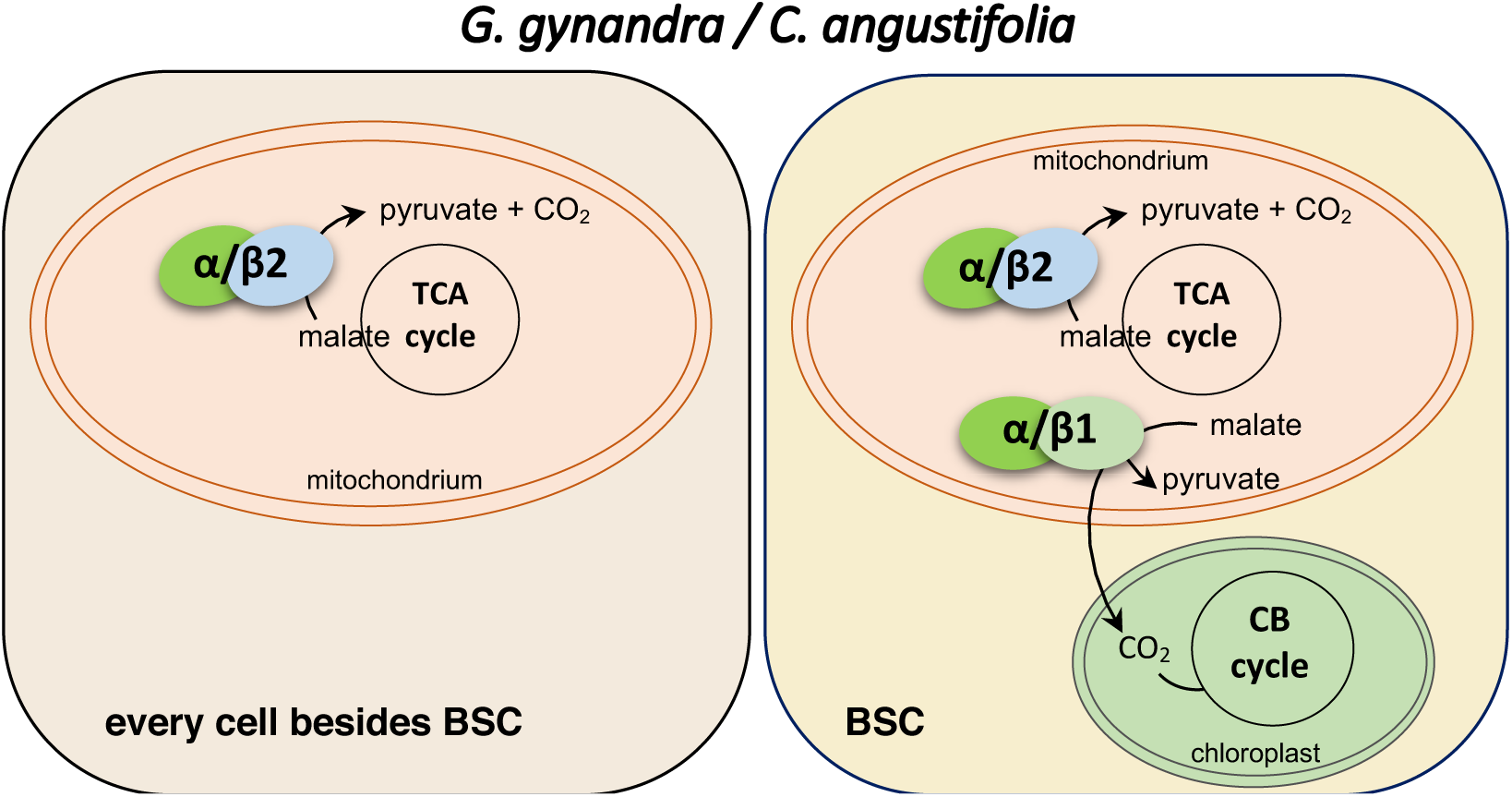
Schematic representation of the localization of C_4_ Cleome *G. gynandra* NAD-ME entities in different types of cells. GgNAD-MEα/β1/β2 is present in mitochondria of every cell besides bundle sheath cells (BSC) where it is involved in malate respiration as an associated enzyme of the Tricarboxylic acid (TCA) cycle. GgNAD-MEα/β1 is present in mitochondria of BSC where it performs the C_4_ decarboxylase activity delivering CO_2_ for the Calvin Benson (CB) cycle.

## DISCUSSION

A straightforward rationale is to assume that if a gene was recruited into the C_4_ pathway, it should be expressed at higher levels than in C_3_ species and its transcript mostly accumulated during the day (Christin et al., 2013b; Moreno-Villena et al., 2018). Our comparative quantitative transcriptional analysis revealed that all *NAD-ME* genes are higher expressed in C_4_ Cleome leaves than their homologs in C_3_ Cleome leaves. Therefore, it is not possible to unambiguously identify a concrete *C_4_-NAD-ME* candidate gene by the expressional analysis, which in part explains why, until to date, a specific C_4_-NAD-ME was never identified at the molecular level.

Here, using a combination of biochemical and proteomic analyses, we identify different NAD-ME isoforms in C_3_ and C_4_ Cleome organs, which arise through a differential combination of subunits previously shown to be selectively adapted through C_4_ evolution (Tronconi et al., 2020). We found that non-photosynthetic and photosynthetic organs of C_3_ Cleome *T. hassleriana* possesses two NAD-ME isoforms, ThNAD-MEα/β1 and ThNAD-MEα/β2, with kinetic properties similar to those of *A. thaliana* NAD-MEs. These results indicate that ThNAD-MEα/β1 and ThNAD-MEα/β2 represent housekeeping enzymes that participate in mitochondrial malate respiration. The paralogs *β1* and *β2* genes were possibly fixed by genetic drift (Tronconi et al., 2020), most probably resulting in redundancy of function for ThNAD-MEα/β1 and ThNAD-MEα/β2 in the C_3_ species. In the C_4_ Cleome species, NAD-MEα/β2 is responsible for the housekeeping function.

We found that NAD-MEα/β1 is exclusively present in leaves of the C_4_ species *G. gynandra* and *C. angustifolia.* Here, NAD-MEα/β1coexist in bundle sheath cells mitochondria with the NAD-ME α/β2 housekeeping heteromer. GgNAD-MEα/β1 has a high affinity for malate and high catalytic efficiency, and its activity is enhanced in the presence of aspartate, an intermediate of the NAD-ME subtype C_4_ pathway. Aspartate is formed in the mesophyll cells from OAA, the product of the first carboxylation step, and is used to shuttle the prefixed CO_2_ to BSC mitochondria (Ludwig, 2016b). In BSC mitochondria, aspartate is converted to malate. In *G. gynandra*, the aspartate aminotransferase recruited into the C_4_ pathway is located in the mitochondria (Sommer et al., 2012). We postulate that aspartate is an excellent candidate to regulate the activity of NAD-ME when functioning as C_4_ decarboxylase. Due to the mentioned kinetic and regulatory properties of GgNAD-MEα/β1 and the fact that this enzymatic entity is only found in leaves of C_4_ Cleome, we conclude that GgNAD-MEα/β1 is the C_4_-NAD-ME photosynthetic isoform.

Our extensive enzymatic analysis indicated that NAD-MEβ1 of both C_4_ species, *G. gynandra* and *C. angustifolia,* are not active. Comparison of closely related β-NAD-ME sequences indicated that NAD-MEβ1 of the C_4_ species accumulated many amino acid changes during evolution (Tronconi et al., 2020). We found that the changes of Y132 and V297, which participate in the catalytic mechanism, are most probably responsible for the catalytic inactivity of Ca/GgNAD-MEβ1. The loss of function of the active site could have led to the generation of an allosteric site for aspartate, a C_4_ compound structurally similar to malate. Our results indicate that aspartate competitively inhibits NAD-MEβ2 (Supplemental Tables 3), suggesting that it binds effectively to the active site. Lui et al. (2002) showed that NAD-ME from *Ascaris suum* catalyzes the oxidative decarboxylation of aspartate to pyruvate and ammonia through a mechanism that likely mimics that of malate. Thus, the evolution from an active to an allosteric site would have required few amino acid substitutions.

We also found that the Q195H substitution in the NAD-MEβ1 has a stabilizing effect in the dimer interface region (Supplemental Figure 7C). The stabilizing influence of NAD-MEβ1 on the associated NAD-MEα most probably enables the differential assembly of the subunits to form independent protein entities acting in different metabolic pathways in the same cell compartment. The flux of malate through these metabolic pathways is likely regulated by the action of aspartate as we found that this C_4_ intermediate highly enhances the efficiency of C_4_-NAD-ME but at the same time decreases the efficiency of the respiratory isoform (Supplemental Tables 3). The contrasting behavior of the NAD-MEβ subunits in C_4_ Cleome points to a specialized role of NAD-MEβ1 in the C_4_ isoform.

Carbon isotopic discrimination and type of Kranz anatomy in *G. gynandra* and *C. angustifolia* indicated that they have two different C_4_ origins (Feodorova et al., 2010; Bayat et al. 2018). Although these C_4_ species display differences in leaf architecture and physiology (Koteyeva et al., 2011), the recruitment of NAD-ME genes into the C_4_ pathway followed a similar molecular mechanism. After fixing the pair of paralogs *β* genes, one copy was co-opted to play a role in C_4_ photosynthesis through parallel amino substitutions. We postulate that the non-catalytic subunit in Cleome C_4_ species, NAD-MEβ1, was evolutionarily fixed because it enabled the formation of a NAD-MEα/β1 heteromer with suitable biochemical properties to fulfill a role in the C_4_ photosynthetic pathway, such as high catalytic efficiency and activation by aspartate.

In summary, our work shows for the first time that a C_4_-exclusive NAD-ME isoform exists, which is formed through the differential assembly of protein subunits that accumulated adaptive mutations during evolution. Our results will now aid in introducing C_4_ traits into C_3_ plants (Ermakova et al., 2020; Lin et al., 2020; von Caemmerer and Furbank, 2016). Ongoing projects can now rationally design the incorporation of a NAD-ME with C_4_ characteristics by introducing the NAD-MEβ1 of C_4_ Cleome or modifying a preexistent extra NAD-MEβ copy.

## METHODS

### Plant growth conditions

*Gynandropsis gynandra* and *Tarenaya hassleriana* were grown under greenhouse conditions in commercially available soil mixed with Cocopor® (Stender, Schermbeck, Germany). *G. gynandra* was germinated in the dark for 3-5 days, while *T. hassleriana* needed approx. 14 days for germination under long day (16h/8h) light regime. Natural light was supplemented with regular filament lamps mounted 1.2 m above the surface ensuring a minimum of 80 µmol m^−2^ s^−1^ PAR. Seedlings were transferred from germination pots to single clay pots after full formation of the cotyledons. Plants were watered from above as needed and single pot soil was premixed with 3 g L^−1^ Osmocote (Scotts Deutschland GmbH, Nordhorn, Germany) as fertilizer. *Arabidopsis thaliana* (ecotype Columbia-0) was grown in soil as described before (Hüdig *et al*., 2015). For immunolocalization studies, plants were grown in 20 L pots, watered daily to avoid drought stress, and fertilized weekly as described by Sage et al. (Sage et al., 2013). Plants were exposed to direct sunlight most days, giving light intensities of 1600 μmol m^−2^ s^−1^.

### Transcriptional analysis of NAD-ME genes

Total RNA from samples were isolated using the RNeasy Mini Kit (Qiagen, Hilden, Germany) from pooled (n = 5) plant leaf material of *G. gynandra* and *T. hassleriana*. Genomic DNA was removed by DNAse treatment (Ambion Inc., Austin, USA), according to the manufacturer’s instructions. cDNA synthesis was performed using RevertAid H Minus Reverse Transcriptase (Thermo Fisher Scientific, Darmstadt, Germany) with oligodT primers according to the manufacturer’s instructions. Primer for quantitative real-time polymerase chain reaction (qRT-PCR) were designed to amplify a PCR product of 130-210 bp length in the region 450-750 bp from the 3’-end of the coding sequence of the mRNA transcript. The respective Actin2 homolog was chosen from nearest orthologs to *A. thaliana* Actin2. Primer efficiency was determined with regards to primer concentration and annealing temperature, and was used to normalize C_t_ values. Primer specificity was ensured using cloned coding sequences from plasmids as test templates and agarose gel visualisation of the respective final qRT-PCR products. qRT-PCR was run on StepOne Plus (Thermo Fisher Scientific, Darmstadt, Germany) equipped with StepOne Software v2.2.2 using KAPA SYBR FAST qPCR Kit Master Mix (2X) ABI Prism (Kapabiosystems, Boston, USA) in a 15 µL reaction. qRT-PCR amplification was performed at: 95 °C 3 min, 40 cycles at 95 °C for 3 s and 65 °C for 20 s, followed by a standard melting curve.

### Cloning

Coding sequences of the NAD-ME subunits of *G. gynandra* and *T. hassleriana* were amplified from leaf cDNA using Phusion Polymerase (Thermo Fisher Scientific, Darmstadt, Germany) and specific primers (Supplemental Table 4). The amplified fragments were sub-cloned into pCR-TOPO-BluntII (Thermo Fisher Scientific, Darmstadt, Germany) and sequenced using EZ-seq sequencing services (Marcorgen Europe, Amsterdam, The Netherlands). The generated TOPO vectors were used as source for generation of cohesive-end insert fragments for restriction site cloning or as templates for further PCR-fragment constructions according to Gibson (Gibson, 2011). The vector pET16b (Merck, Darmstadt, Germany) was used for heterologous expression in *E. coli*. The pET16 constructions produce NAD-MEs containing a His•Tag (10 histidine residues) sequence fused to the N-terminus for its purification through affinity chromatography. When using classic restriction enzyme cloning, the pET16b vector was linearized with restriction enzymes (*Nde*I/*Xho*I/*BamH*I) as well as the respective PCR products to generate the cohesive ends and circular plasmids were reconstituted with T4 Ligase (Thermo Fisher Scientific, Darmstadt, Germany). If Gibson assembly was used, the primers were designed with a 20 bp overlap with the destination vector. Purified Gibson-assembly PCR-fragments of NAD-ME coding sequences were cloned into pET16b linearized with *BamH*I. Gibson isothermal assembly was performed according to Gibson (Gibson, 2011) with reagents received from Thermo Fisher Scientific (Darmstadt, Germany) and New England Biolabs (Ipswich, USA). Final plasmids were sequenced and amplified using *E. coli* DH5α cells (Thermo Fisher Scientific, Darmstadt, Germany).

For the co-expression of NAD-ME subunits, the cDNA fragments corresponding to the mature NAD-MEα subunits of *G. gynandra* and *T. hassleriana* were cloned in the pET32 vector (Novagen) using specific restriction sites. The restriction sites *NcoI*/*SalI* were used for GgNAD-MEα and *BamHI*/*SalI* for GgNAD-MEβ1 and -β2. The pET32-NAD-MEα constructions produce NAD-MEα containing Trx•Tag (109 aa of thioredoxin) and His•Tag sequences fused to the N-terminus of NAD-ME. On the other hand, the cDNA fragments coding for the mature NAD-MEβ subunits were cloned in the pET29 vector (Novagen), which adds a C-terminal His•Tag sequence. The pET29-NAD-MEβ constructions express the mature NAD-MEβs without the fusion vector-coded sequence because the cloned cDNA has a stop codon in their 5’end. For the co-expression of GgNAD-MEβ1 and -β2, alternatively one β subunit-coding sequence was cloned in the pET32 vector.

The coding sequence of the mature *C. angustifolia* NAD-MEβ1 subunit was synthetized using the BioCat commercial gene synthesis service (Heidelberg, Germany) and cloned into pET32b via *NcoI/BamHI* restriction sites. The resulting plasmid was sequenced and verified using the Marcogen sequencing service.

### Heterologous expression and purification of recombinant NAD-MEs

Production of recombinant single NAD-ME subunits was performed with the T7-polyperase IPTG-inducible expression system. For production of His-tagged GgNAD-MEα and ThNAD-MEα we used *E. coli* BL21 (DE3) cells (Merck, Darmstadt, Germany) (from now on denoted as protocol 1, P1) and for production of His-tagged GgNAD-MEβ1, GgNAD-MEβ2, CaNAD-MEβ1, ThNAD-MEβ1, and ThNAD-MEβ2 we used *E. coli* ArcticExpress (DE3) cells (Agilent technologies, Santa Clara, USA) (from now on denoted as protocol 2, P2). Chemically competent *E. coli* cells were transformed with 50 ng of the expression vector and grown on LB agar plates with antibiotics according to the manufacturer instructions.

For the co-expression of NAD-ME subunits, *E. coli* BL21 (DE3) cells were simultaneously transformed with the pET29 and pET32 constructions. The cells were selected on LB-agar plates supplemented with 100 µg/ml ampicillin (pET32 selection agent) and 50 µg/ml kanamycin (pET29 selection agent). For protein expression, 800 ml liquid cell cultures containing 100 µg/mL ampicillin and 50 µg/mL kanamycin were inoculated with freshly grown over-night culture for 3 h to an OD_600_ of 0.6. Protein expression was induced with 1 mM IPTG. Cells were harvested after 16 h at 16° C by centrifugation at 4° C and 6000 g, and stored at - 20 °C until use. The interspecies chimera AtNAD-MEα/GgNAD-MEβ1 was obtained by cotransformation of *E. coli* BL21 (DE3) with pET32-AtNAD-MEα (Tronconi et al., 2008) and pET29-GgNAD-MEβ1.

For the induction of protein expression and purification, see the Supplementary information.

### Protein quantification and gel electrophoresis procedures

Sodium dodecyl sulphate polyacrylamide gel electrophoresis (SDS-PAGE) was used to monitor protein purification according to Laemmli (1970). Gradient native PAGE (Novex 4-12% Tris-Glycine gels, Invitrogen, Thermo Fisher Scientific) was used to separate native protein complexes from plant tissue by omitting β-mercaptoethanol and SDS from any buffer used for sample loading, pouring or running PAGE gels. Gels were run at 4°C and 100V. Proteins in gels were visualized using Coomassie brilliant blue staining procedure. Protein concentration was determined by using BioRad Protein Assay (BioRad, Hercules, United States) or alternatively, a simplified amido black 10B precipitation method (Schaffner and Weissmann, 1973) using total serum protein as standard.

### Preparation of protein extracts for native PAGE and *in-gel* NAD-ME activity assays

Plant tissue was freshly harvested from greenhouse grown *G. gynandra*, *T. hassleriana*, *C. angustifolia*, and *A. thaliana* and ground to fine powder in liquid nitrogen. Extraction of soluble proteins was performed by adding 3 µL per mg fresh weight ice-cold extraction buffer (0.1 M Tris-HCl pH 8.0, 0.1 M NaCl, 0.5 % (v/v) Triton X-100, 2 mM PMSF, 10 mM dithiothreitol (DTT), 0.1% (w/v) polyvinylpyrrolidone 40), followed by vigorous vortexing and incubation for 20 min on ice. Soluble protein was separated by centrifugation (10000 g, 10 min, 4° C) and supernatant was transferred to a new tube. Proteins were separated in native PAGE run at 100 V at 4° C.

After electrophoresis, gels were assayed for NAD(P)-ME activity. For NAD-ME and NADP-ME activity assay, gels were first incubated in 50 mM MOPS-NaOH (pH 6.8) at room temperature for 30 min. Rebuffered native PAGE gels were incubated in 50 mM MOPS-NaOH (pH 6.8), 0.05 % (w/v) nitro blue tetrazolium (NBT), 150 μM PMS, 10 mM MnCl_2_, 5 mM NAD or 5 mM NADP, and either 20 mM, 100 mM or no malate. Gels were documented after initial clearance in ddH_2_O and gel slices with NAD-ME activity were processed for protein identification by mass spectrometry.

### Mass spectrometry

Slices of PAGE samples for mass spectrometry were essentially processed as described previously (Brenig et al., 2020). Briefly, protein containing bands were washed, reduced by dithiothreitol, alkylated with iodoacetamide and digested with trypsin. Resulting peptides were extracted from the gel and cleaned up using solid phase extraction (HLB µElution plate, Waters) and finally resuspended in 0.1% trifluoroacetic acid and analyzed by liquid chromatography coupled mass spectrometry. Peptide separation was carried out on an Ultimate 3000 rapid separation system (Thermo Fisher Scientific) on 25 cm long C18 columns using a 54 min separation gradient (2 h for *G. gynandra* leaf band migration level 1 sample 2) essentially as described in (Poschmann et al., 2014). Subsequently, separated peptide were injected via a nano-electrospray interface into the mass spectrometer operated in data dependent positive mode and searches were carried out in the proteome discoverer framework as described in the supplemental information.

### Kinetic analysis

Purified recombinant NAD-MEs were assayed for the oxidative decarboxylation of malate using a standard reaction mixture containing 50 mM MES-NaOH pH 6.2-6.4, 10 mM MnCl_2_, 4 mM NAD, 20 mM malate in a final volume of 0.5 ml. The reaction was started by the addition of malate. An extinction coefficient (*ε*) of 6.22 mM^−1^ cm^−1^ for NADH at 340 nm was used in the calculations. Initial-velocity studies were performed by varying the concentration of one of the substrates around its *K*_m_ value while keeping the other substrate concentrations at subsaturating or saturating levels. All kinetic parameters were calculated at least by triplicate determinations and adjusted to non-lineal regression using free concentrations of all substrates.

One unit (U) is defined as the amount of enzyme that catalyzes the formation of 1 mmol of NADPH per min under the specified conditions.

When testing the ability of metabolic intermediates as possible modulators (inhibitors or activators) of the enzymatic activity, NAD-ME activity (1-3 ug of protein) was measured in the absence or presence of 0.5 or 5.0 mM of this metabolite, at a saturating NAD concentration (4 mM) and sub-saturating levels of malate (1 mM for the housekeeping isoforms and 0.1 mM for GgNAD-MEα/β1), which were kept at the one-third of the Km value for each enzyme (Km/3). The apparent activation constant (A_50_) values were obtained by varying the concentration of activator, between 0.1 and 30 mM, while keeping the NAD concentrations at a saturating level (4 mM) and malate at non-saturating concentrations (∼3 mM for the housekeeping isoforms and ∼ 0.3 mM for GgNAD-MEα/β1).

### Gel Filtration Chromatography

Molecular masses of recombinant GgNAD-MEs were evaluated by gel-filtration chromatography on an ÄKTA purifier system (GE Healthcare) using a Biosep-SEC-S3000 column (Phenomenex). The column was equilibrated with 50 mM MES-NaOH, pH 6.5, and calibrated using molecular mass standards. and calibrated using molecular mass standards (Sigma-Aldrich, Merck, Darmstadt, Germany). The samples and the standards were applied in a final volume of 100 µL at a constant flow rate of 0.5 mL/min. All the experiments were performed in triplicate.

### Generation of comparative models

We used SWISS-MODEL (Waterhouse et al., 2018) to build comparative models of the α, β1, and β2 isoforms of GgNAD-ME as well as the α and β_1_ isoform of ThNAD-ME. All models were built using PDB ID 1PJ3 (Tao et al., 2003) as a template structure (sequence identities: 42.0-42.8 %, coverage: > 88 %). The structures of the monomeric isoforms were used to build the GgNAD-MEα/α homodimer as well as the GgNAD-MEα/β1, GgNAD-MEα/β2, and ThNAD-MEα/β1 heterodimers by superimposing the monomers onto PDB ID 1PJ3. For further details, see the Supplemenatry information.

### Molecular dynamics simulations

For details on the preparation of the molecular systems for MD simulations, see the Supplementary information. All MD simulations were performed using the graphics processing unit version of pmemd (Le Grand et al., 2013; Salomon-Ferrer et al., 2013). The Langevin thermostat (Pastor et al., 1988) was used for temperature control with a collision frequency of 2 ps^−1^. The particle mesh Ewald method (Berendsen et al., 1984) was used for handling long-range interactions with a cutoff of 8.0 Å. The SHAKE algorithm (Ryckaert et al., 1977) was used to constrain bonds to hydrogens. The simulations were performed with an integration time step of 2 fs. For details on system minimization and thermalization, see the Supplementary information.

For production, the thermalized systems were simulated for a further 500 ns of NPT-MD without any restraints. For each system, five separate thermalizations and production simulations were performed, yielding an aggregate production simulation time per system of 2.5 ms. CPPTRAJ (Roe and Cheatham, 2013) was used to analyze the trajectories. Visualization of the molecular structures was done with PyMOL (DeLano, 2002).

### Constrained Network Analysis

Rigidity analysis was performed using the CNA software package (Pfleger et al., 2013). CNA is a front- and back-end to FIRST (Jacobs et al., 2001; Pfleger et al., 2013), which is used to construct networks of covalent and non-covalent interactions from input structures (Hermans et al., 2017). The strength of hydrogen bonds and ionic interactions are calculated by FIRST. Hydrophobic interactions are considered if the van der Waals spheres of two atoms are less than 0.25 Å apart. By successively removing the constraints below a certain energy cutoff *E_cut_* (−7.0 kcal mol^−1^ ≤ *E_cut_* < −0.1 kcal mol^−1^, Δ*E_cut_* = 0.1 kcal mol^−1^) from the network topologies, “constraint dilution trajectories” are generated (Radestock and Gohlke, 2008; Radestock and Gohlke, 2011). For details on how changes of the biomolecular stability along the “constraint dilution trajectories” are quantified (Rathi et al., 2015; Wifling et al., 2019) and how the impact of individual substitutions in the GgNAD-MEβ1 isoform on the biomolecular rigidity and flexibility was assessed using a perturbation approach (Pfleger et al., 2017), see the Supplementary information.

### Immunolocalization studies

Leaf tissue sampled from the middle of the youngest fully expanded leaf and root tissue sampled from the mature region of the youngest roots were fixed in 0.5 % glutaraldehyde, 4 % paraformaldehyde in sodium cacodylate buffer and prepared for transmission electron microscopy (TEM) and immunohistochemistry as described by Khoshravesh et al. (Khoshravesh et al., 2017). Specific antibodies were raised against GgNAD-ME subunits based on small unique peptide sequences as antigens in rabbits. GgNAD-MEα and GgNAD-MEβ2 antibodies were produced by Agrisera (Vännäs, Sweden), GgNAD-MEβ1 antibodies were produced by Cambridge Research Biochemicals (Billingham, UK). The dilutions used were 1:100 (GgNAD-MEα), 1:50 (GgNAD-MEβ1, GgNAD-MEβ2), and 1:20 (secondary antibody; 18-nm colloidal gold-affiPure goat anti-rabbit IgG; Jackson Immunoresearch). Incubation times in the primary and secondary antibodies were 2 and 1 h, respectively. Controls were run by omitting primary antibody. TEM images were captured on a Phillips 201 transmission electron microscope equipped with an Advantage HR camera system (Advanced Microscopy Techniques, Woburn, MA, USA).

### Statistical analyses

Statistical analysis for all experiments was performed using GraphPad Prism (v. 6.01) software using a two-way analysis of variance (ANOVA; with α<0.05) and Tukey correction for post-hoc multiple comparisons.

## Data availability

The mass spectrometry proteomics data have been deposited to the ProteomeXchange Consortium via the PRIDE partner repository with the dataset identifier PXD026547 and 10.6019/PXD026547. The data supporting the findings of this manuscript are available from the corresponding author upon reasonable request.

## Corresponding author

Correspondence and request for material should be requested to Veronica G. Maurino.

## ACKNOWLEDGEMENTS

This work was supported by grants of the Deutsche Forschungsgemeinschaft MA2379/18-1 and the Germany’s Excellence Strategy EXC1028 to VGM, and the Canadian Natural Science and Engineering Research Council grant 2015-04878 to TLS. HG is grateful for computational support by the “Zentrum für Informations und Medientechnologie” at the Heinrich-Heine-Universität Düsseldorf and the computing time provided by the John von Neumann Institute for Computing (NIC) on the supercomputer JUWELS at Jülich Supercomputing Centre (JSC) (user IDs: HKF7, VSK33).

The authors have no competing interests (financial/non-financial) that might be perceived to influence the interpretation of the article.

## AUTHOR CONTRIBUTIONS

Conceptualization and Supervision, VGM and MAT. Investigation, MH, MAT, JPS, TLS, MS, GP, and DB. Data curation GP. Formal Analysis, HG and DB. Writing – all authors. Funding Acquisition, VGM and TLS.

## SUPPLEMENTAL FIGURES AND TABLES

**Supplemental Figure 1.**
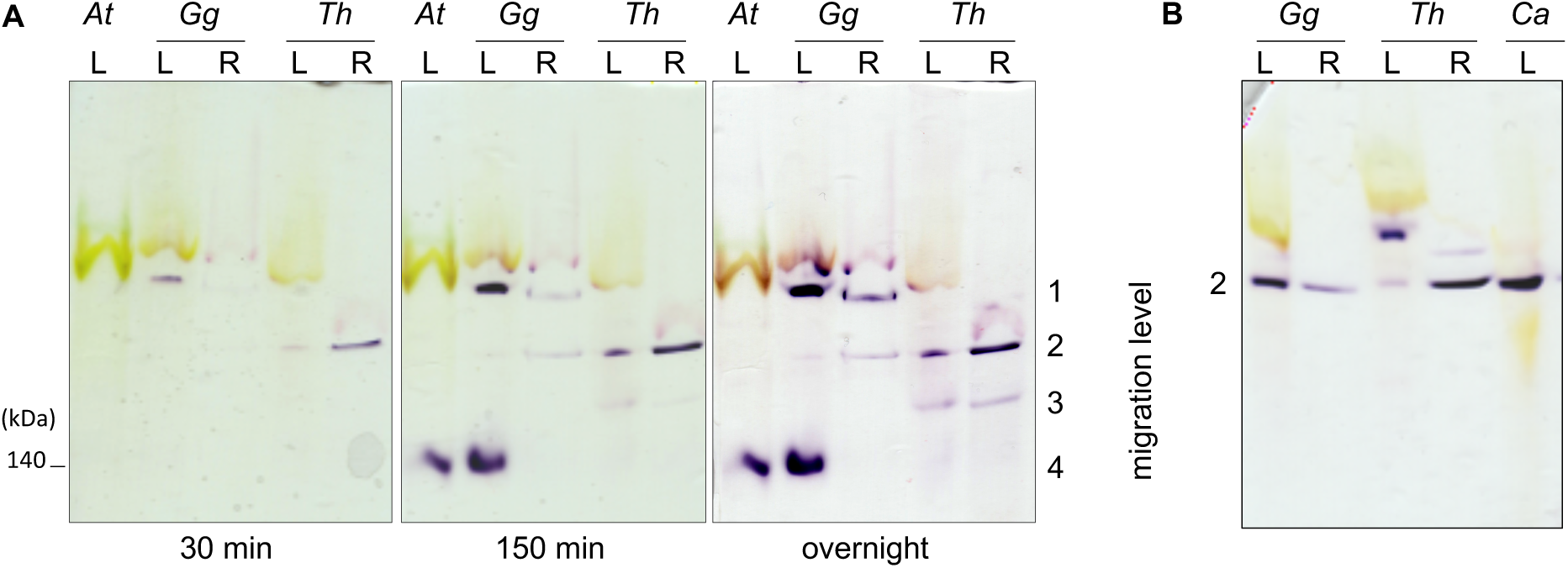
Behavior of NAD-ME entities of Cleome species. (A) Gradient native PAGE of soluble protein extracts of leaves (L, 40 µg protein) and roots (R, 20 µg protein) of *G. gynandra* (Gg), and *T. hassleriana* (Th) incubated in the assay medium with NAD as cofactor at pH 6.8. Incubation times are given below the gels. (B) Gradient native PAGE of soluble protein extracts of leaves (L, 40 µg protein) and roots (R, 20 µg protein) of *G. gynandra* (Gg), *T. hassleriana* (Th), and *C. angustifolia* (Ca) were incubated overnight in the assay medium with NADP as cofactor at pH 6.8. A violet precipitate indicates NAD(P)-ME activity. Migration levels 1 to 4 are indicated on the right. The molecular weight of AtNAD-ME (Tronconi et al., 2008) is indicated on the left.

**Supplemental Figure 2.**
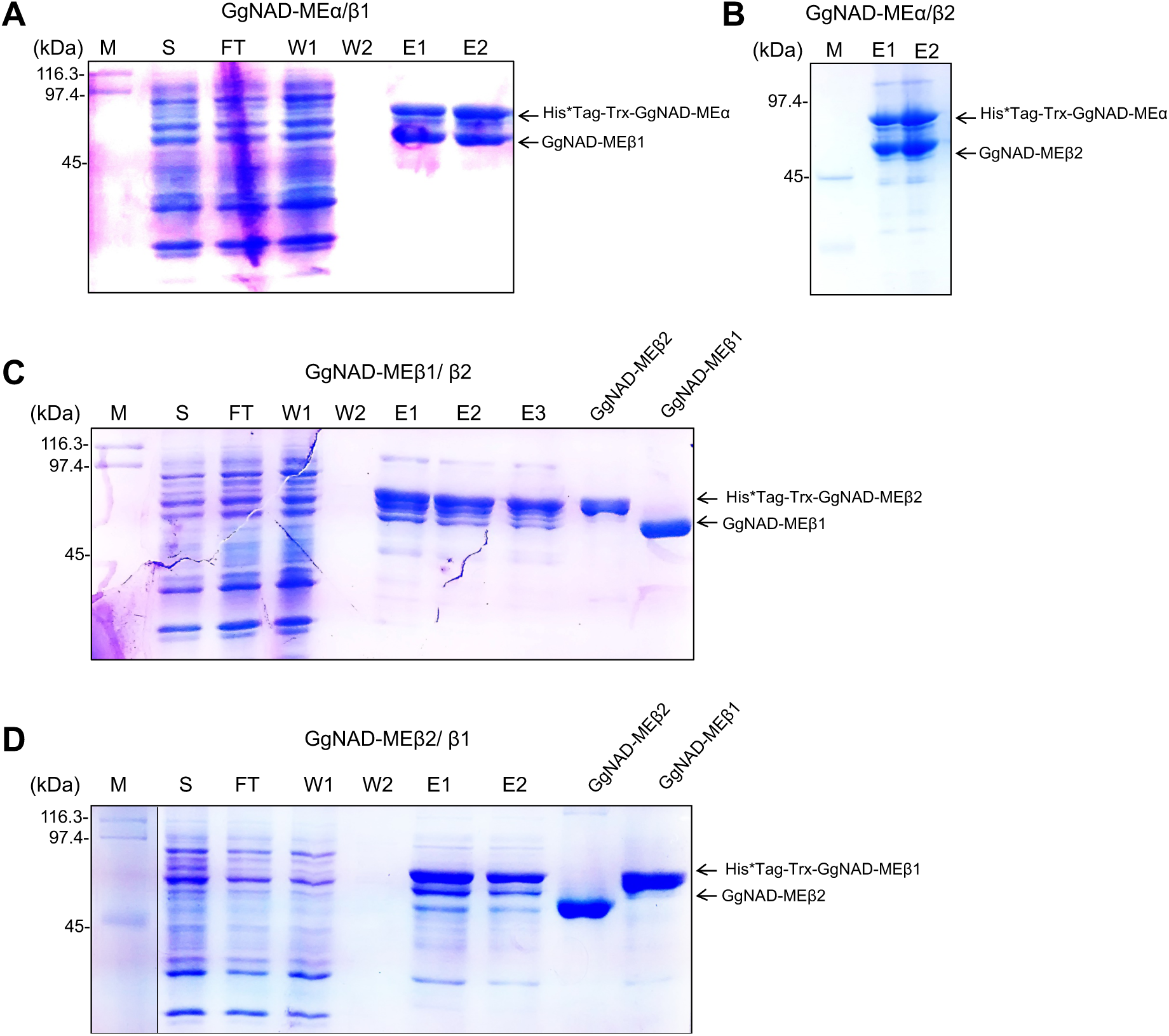
Representative SDS-PAGEs showing purification steps of recombinant NAD-ME from *E. coli* BL21 cells transformed with: (A) pET32-GgNAD-MEα and pET29-GgNAD-MEβ1 to produce GgNAD-MEα/β1; (B) pET32-GgNAD-MEα and pET29-GgNAD-MEβ2 to produce GgNAD-MEα/β2; (C) pET32-GgNAD-MEβ2 and pET29-GgNAD-MEβ1, and (D) pET32-GgNAD-MEβ1 and pET29-GgNAD-MEβ2 to analyse the interaction between β-subunits. M, molecular weight markers; S, supernatant containing soluble protein after sonication; FT, flow through; W1-2, washing fractions with increasing imidazole concentrations; E1-3, elution fractions. For simplicity, only elution fractions are shown in (B). His*tag-Trx: 109 amino acid residues corresponding to thioredoxin and the His*Tag are fused to the N-terminal end of GgNAD-MEs.

**Supplemental Figure 3.**
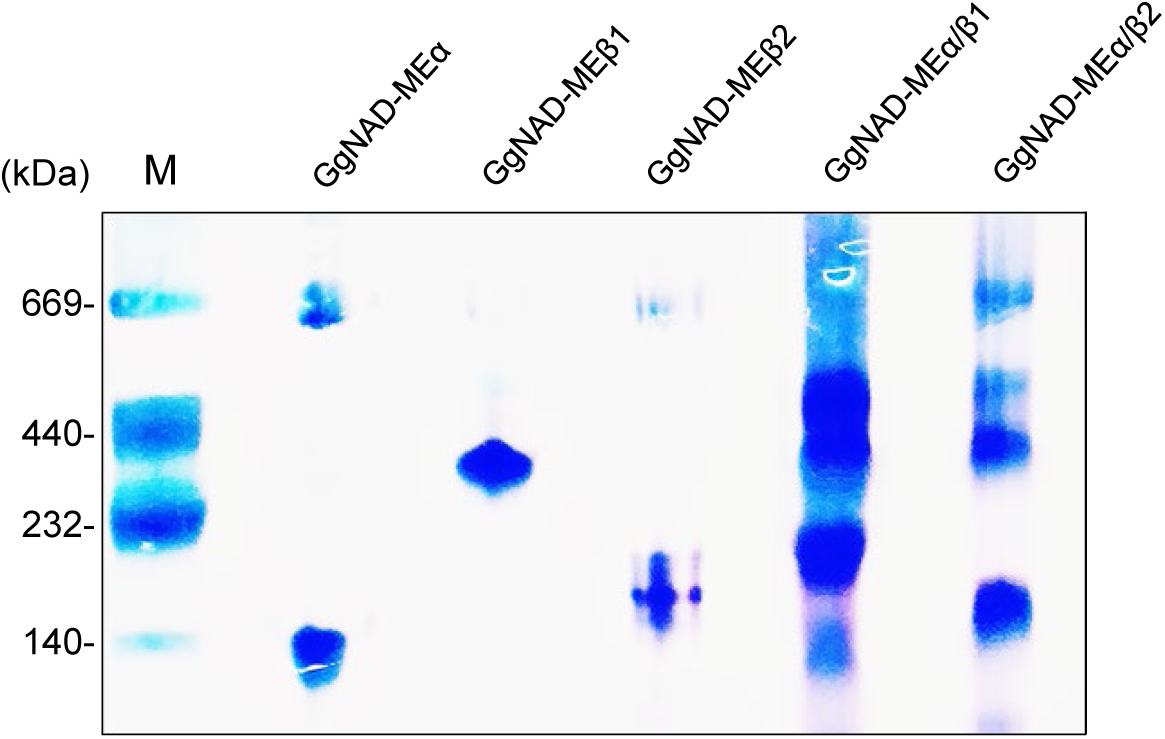
Coomassie-stained native PAGE of purified recombinant GgNAD-MEs. GgNAD-MEα, GgNAD-MEβ1and GgNAD-MEβ2 (10 µg protein), and GgNAD-MEα/β1 and GgNAD-MEα/β2 (20 µg protein) were loaded in each case. M, molecular weight markers.

**Supplemental Figure 4.**
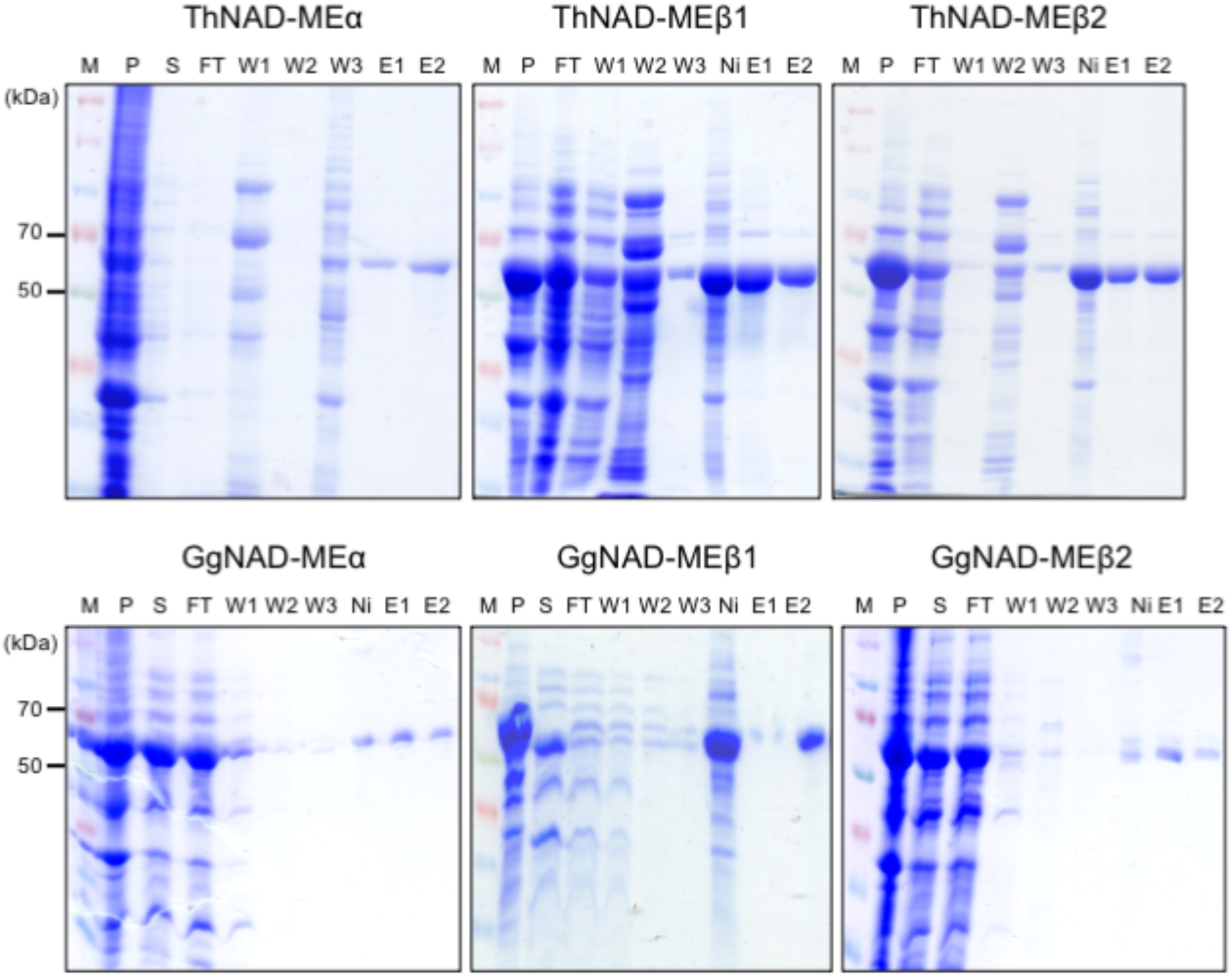
Representative SDS-PAGEs showing different purification steps of the recombinant NAD-ME single proteins of *T. hassleriana* and *G. gynandra*. Expected molecular weights ∼67 kDa: ∼64 kDa of mature protein plus ∼2.5 kDa of the His-Tag or Trx-His-Tag ∼17kDa). M, molecular weight marker; P, cell debris pellet after sonication; S, supernatant containing soluble protein after sonication; FT, flow through; W1-3, washing fractions with increasing imidazole concentrations; Ni, proteins bound to Ni-NTA agarose after elution; E1-2, elution fractions.

**Supplemental Figure 5.**
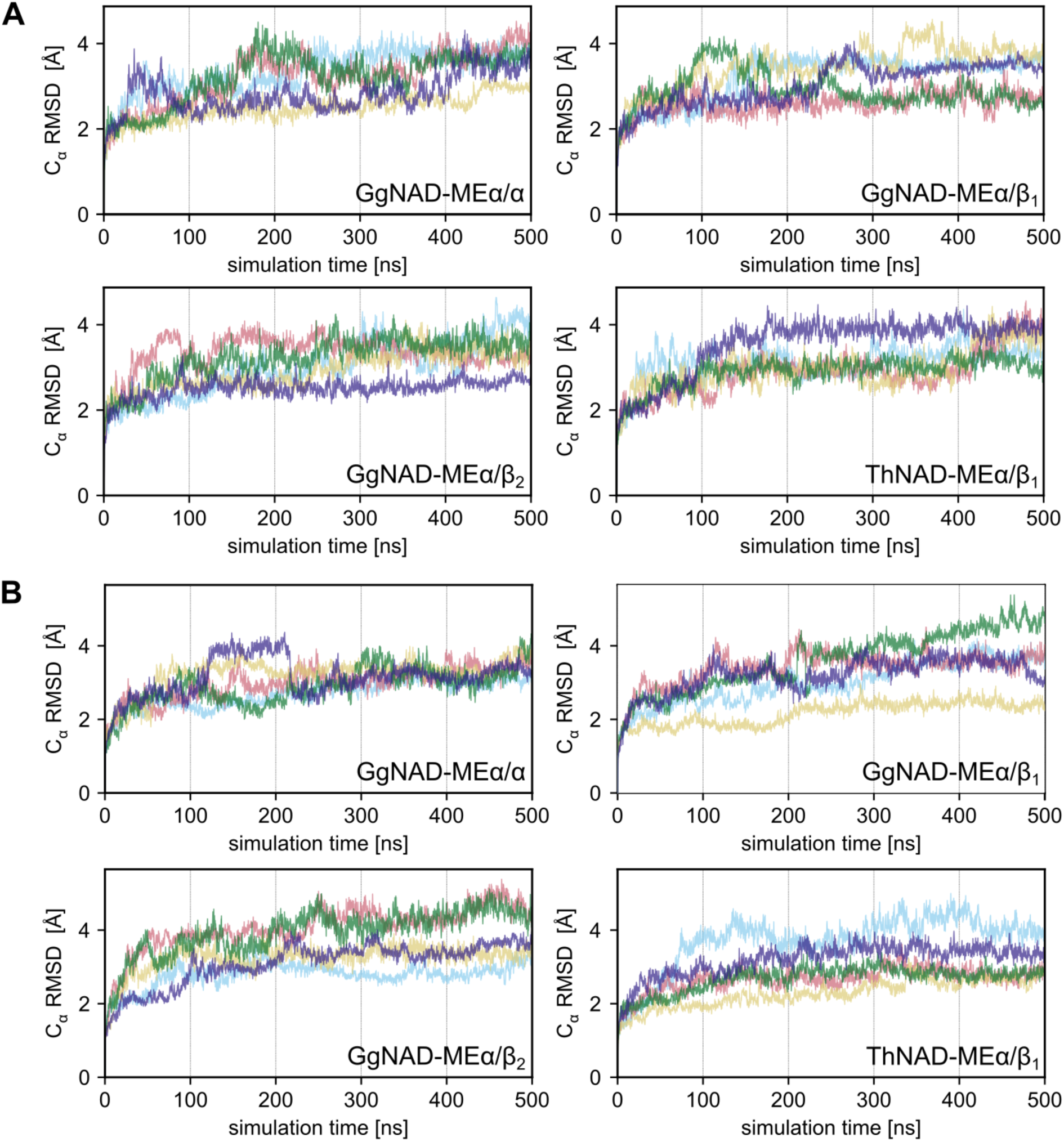
Structural stability of the simulated systems. For the four systems and five independent MD simulations each, the C_α_ RMSD of the “catalytic” subunit (A) and the “effector” subunit (B) are depicted as a function of the simulation time. Despite the considerable sizes of the systems, the RMSD of the subunits generally plateaus at < 4 Å after ∼200 ns of simulation time.

**Supplemental Figure 6.**
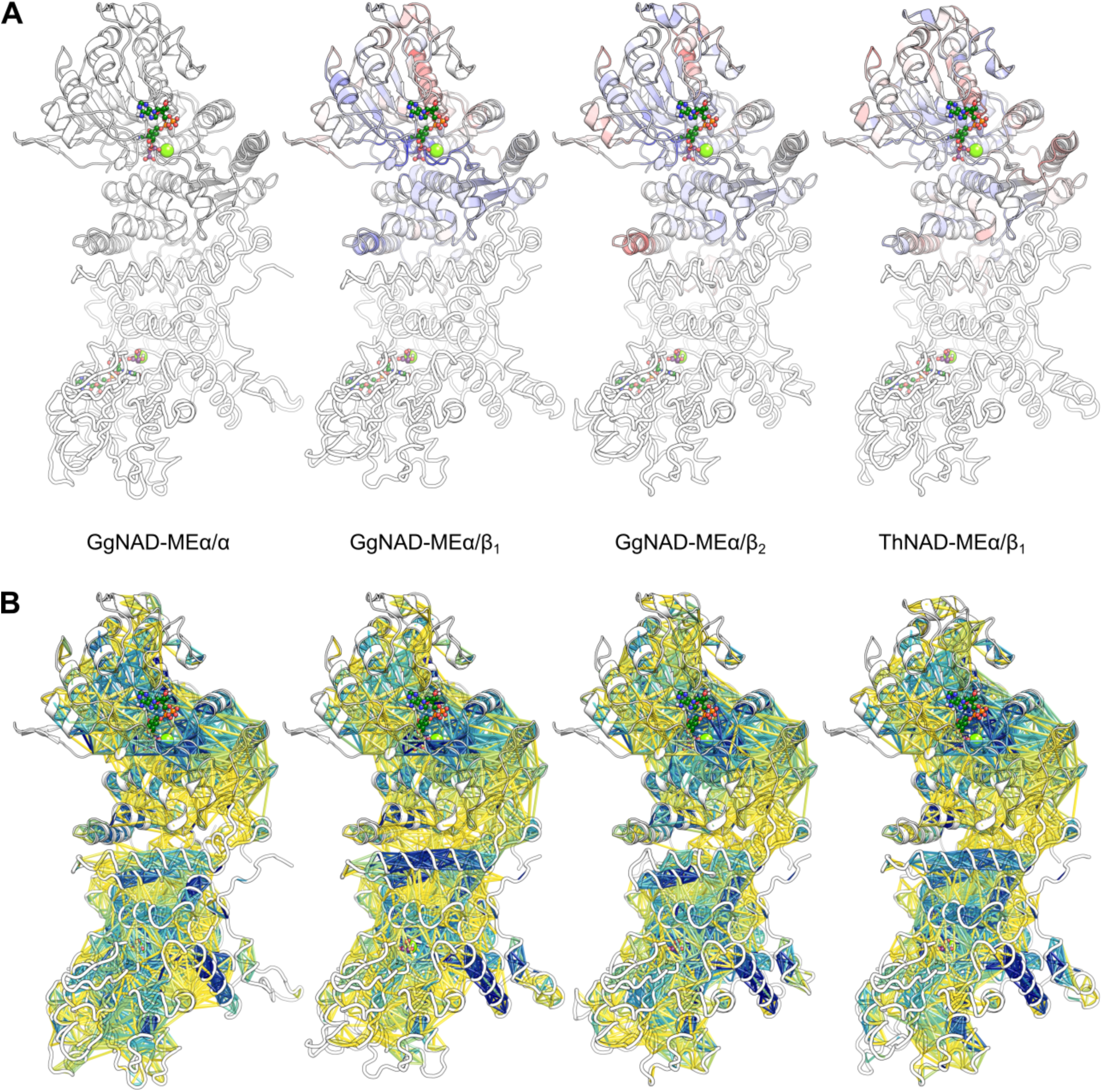
Constraint Network Analysis. (A) Structures of the dimeric complexes of the NAD-ME with bound NAD^+^ and L-malate. The “catalytic” subunit (depicted as helices and sheets) is shown at the top. The effector subunit is shown as ribbons below. The coloring corresponds to the differences in the chemical potential energy per residue (Δ*E*_CNA,*i*_) with respect to the “catalytic” subunit of the GgNAD-MEα/α homodimer, which is therefore not colored. Red (blue) colors indicate regions that are less (more) structurally rigid than in GgNAD-MEα/α. (B) Depiction of rigid contact strength between amino acid pairs in the homodimer and heterodimers. A rigid contact denotes that two residues are part of a structurally rigid cluster in the structure. Blue (yellow) colors indicate more (less) prevalent rigid contacts. The “catalytic” α subunit is depicted as helices and sheets at the top, the “effector” subunit as ribbons below. In the GgNAD-MEα/β1 heterodimer, there are more rigid contacts across the interface than in the other three dimers.

**Supplemental Figure 7.**
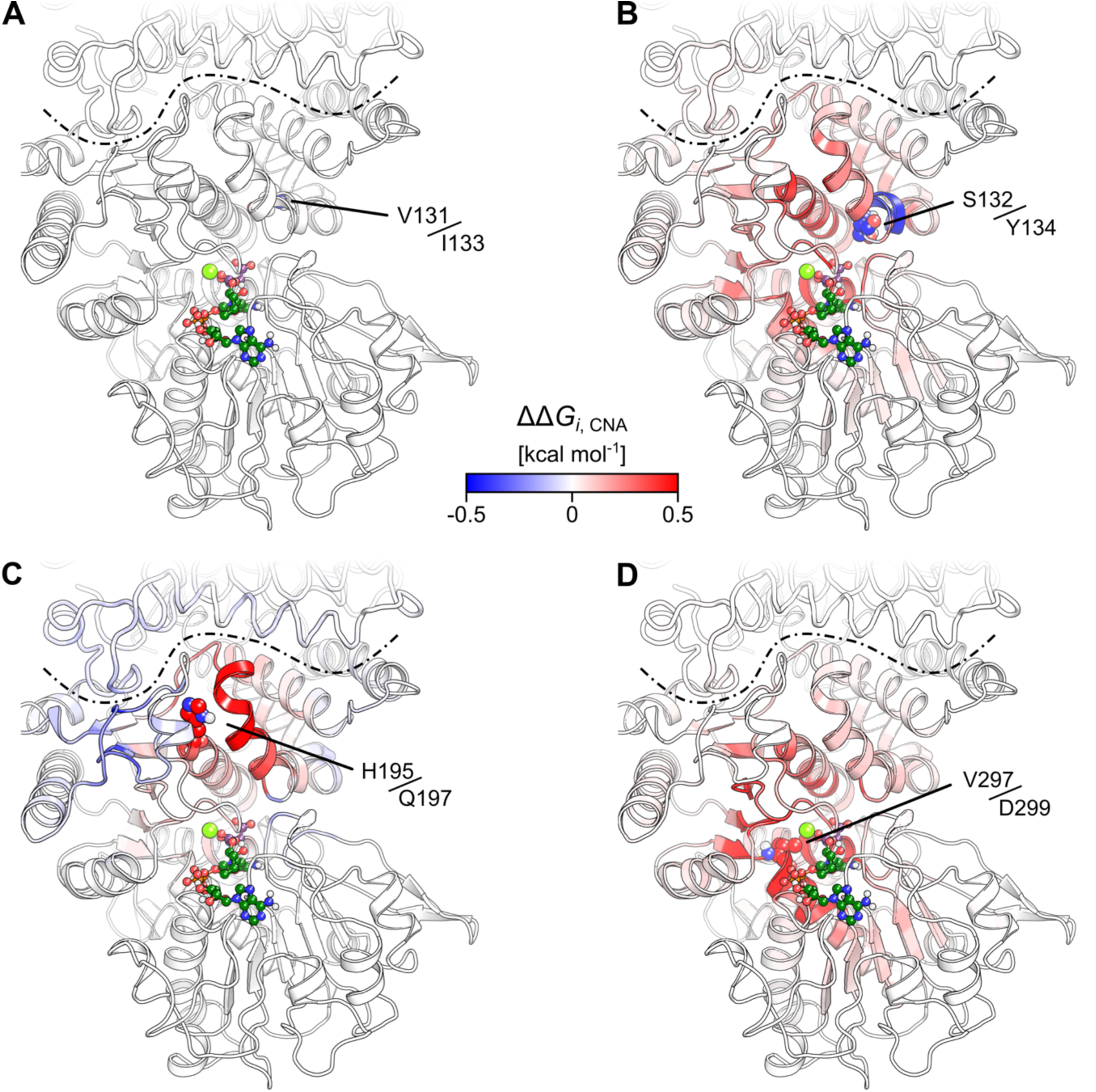
Impact of the substitutions in GgNAD-MEβ_1_ on the structural rigidity. The per-residue difference in structural stability and rigidity (ΔΔ*G_i_*_,CNA_) due to the indicated substitution is mapped on the structure of GgNAD-MEα/β1. The separation between the α-subunit (shown as ribbon, top) and the β-subunit (shown as sheets and helices, bottom) is shown by a dashed black line. For each substitution, the location is indicated in the structure (spheres). The corresponding label shows the type and sequence position of the residue in GgNAD-MEβ1 (A and B) and GgNAD-MEβ2 (C and D). The color scale applies to all depicted structures; blue (red) colors indicate increased (decreased) structural rigidity when perturbing the GgNAD-MEβ1 residue *versus* the GgNAD-MEβ2 one (eq. 2).

**Supplemental Figure 8.**
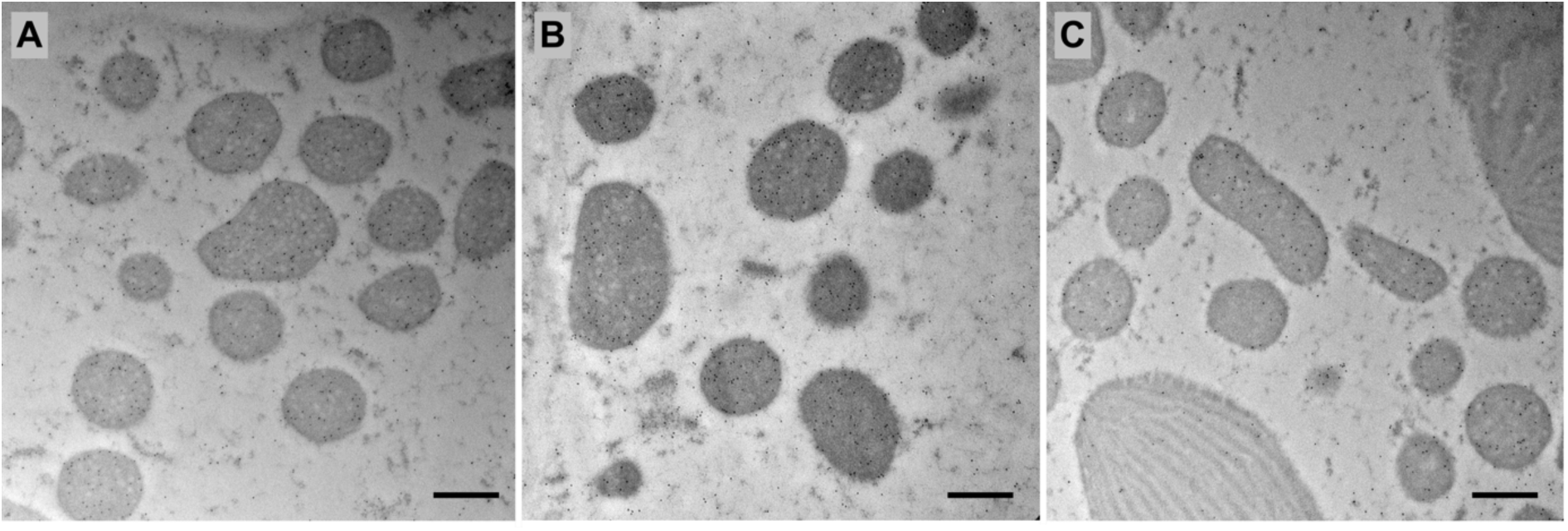
Transmission electron micrographs of bundle sheath cells of *Gynandropsis gynandra* showing mitochondria with immunogold labelling (black particles) of GgNAD-MEα (A), GgNAD-MEβ1 (B), and GgNAD-MEβ2 (C). Bars, 500 nm.

**Supplemental Table 1.**
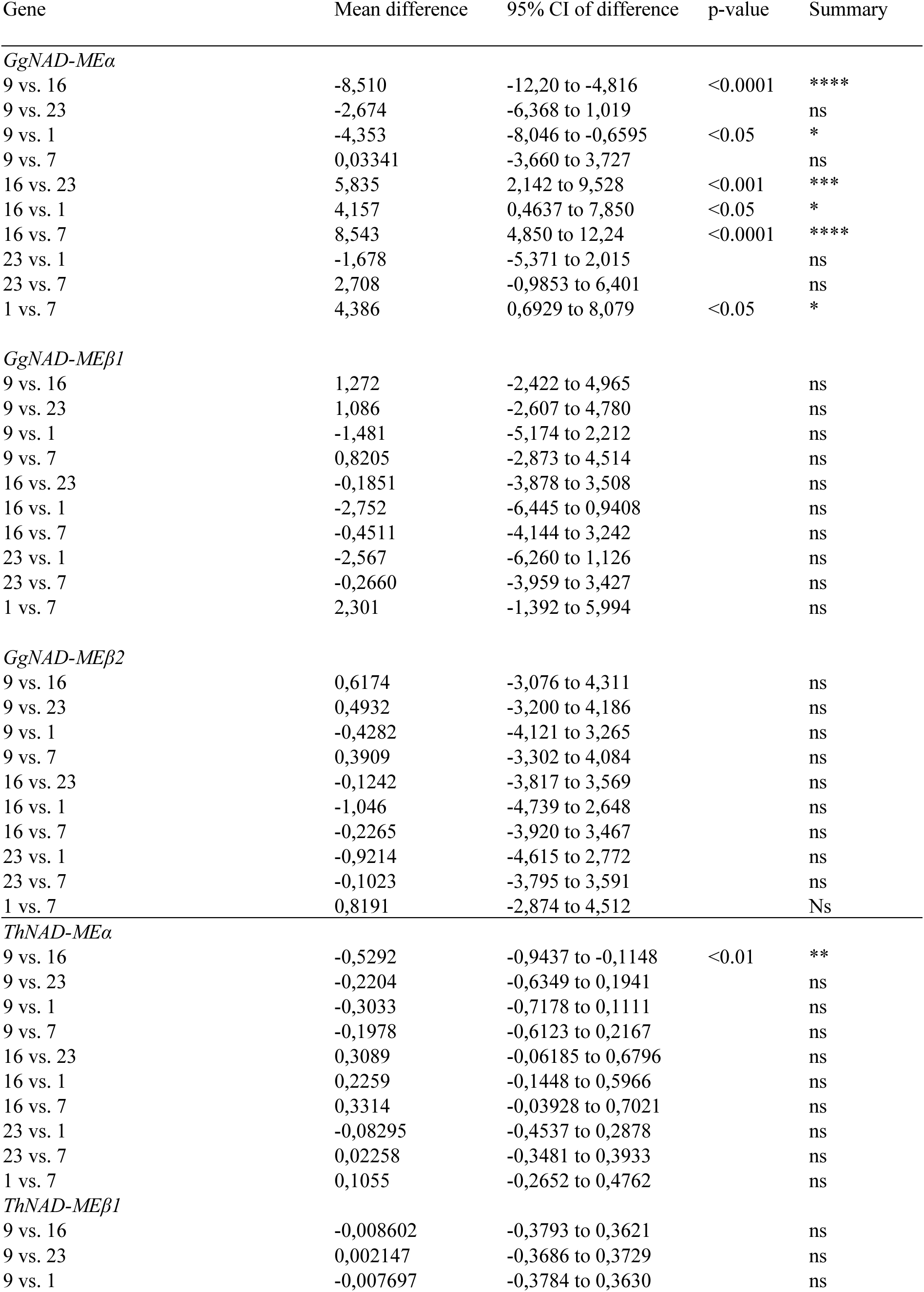
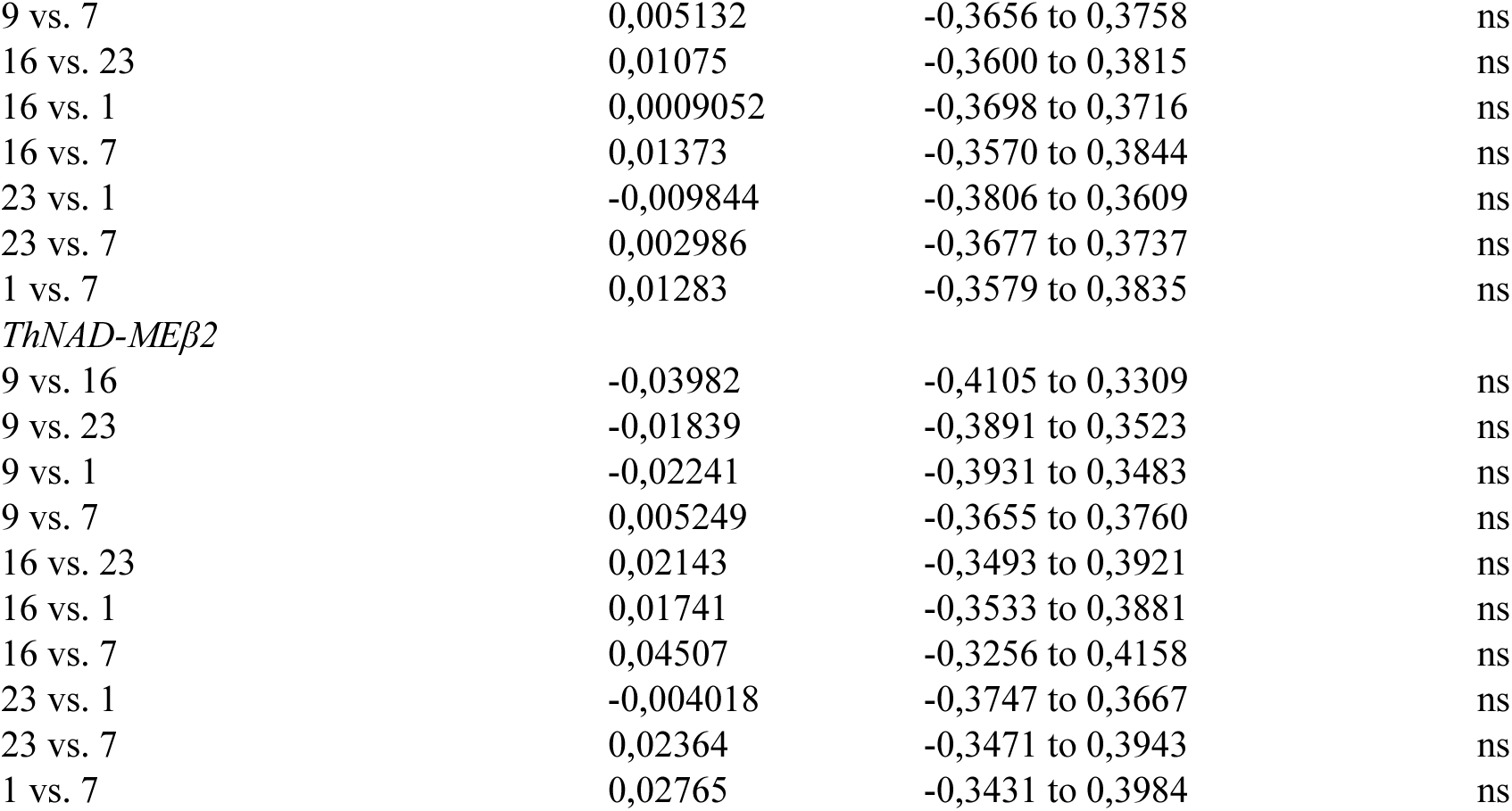
Summary of the Tukey’s multiple comparisons test for data of Figure 1. CI=confidence interval. NS=not significant.

| Gene | Mean difference | 95% CI of difference | p-value | Summary |
| --- | --- | --- | --- | --- |
| *GgNAD-MEα* |  |  |  |  |
| 9 vs. 16 | -8,510 | -12,20 to -4,816 | <0.0001 | **** |
| 9 vs. 23 | -2,674 | -6,368 to 1,019 |  | ns |
| 9 vs. 1 | -4,353 | -8,046 to -0,6595 | <0.05 | * |
| 9 vs. 7 | 0,03341 | -3,660 to 3,727 |  | ns |
| 16 vs. 23 | 5,835 | 2,142 to 9,528 | <0.001 | *** |
| 16 vs. 1 | 4,157 | 0,4637 to 7,850 | <0.05 | * |
| 16 vs. 7 | 8,543 | 4,850 to 12,24 | <0.0001 | **** |
| 23 vs. 1 | -1,678 | -5,371 to 2,015 |  | ns |
| 23 vs. 7 | 2,708 | -0,9853 to 6,401 |  | ns |
| 1 vs. 7 | 4,386 | 0,6929 to 8,079 | <0.05 | * |
| *GgNAD-MEβ1* |  |  |  |  |
| 9 vs. 16 | 1,272 | -2,422 to 4,965 |  | ns |
| 9 vs. 23 | 1,086 | -2,607 to 4,780 |  | ns |
| 9 vs. 1 | -1,481 | -5,174 to 2,212 |  | ns |
| 9 vs. 7 | 0,8205 | -2,873 to 4,514 |  | ns |
| 16 vs. 23 | -0,1851 | -3,878 to 3,508 |  | ns |
| 16 vs. 1 | -2,752 | -6,445 to 0,9408 |  | ns |
| 16 vs. 7 | -0,4511 | -4,144 to 3,242 |  | ns |
| 23 vs. 1 | -2,567 | -6,260 to 1,126 |  | ns |
| 23 vs. 7 | -0,2660 | -3,959 to 3,427 |  | ns |
| 1 vs. 7 | 2,301 | -1,392 to 5,994 |  | ns |
| *GgNAD-MEβ2* |  |  |  |  |
| 9 vs. 16 | 0,6174 | -3,076 to 4,311 |  | ns |
| 9 vs. 23 | 0,4932 | -3,200 to 4,186 |  | ns |
| 9 vs. 1 | -0,4282 | -4,121 to 3,265 |  | ns |
| 9 vs. 7 | 0,3909 | -3,302 to 4,084 |  | ns |
| 16 vs. 23 | -0,1242 | -3,817 to 3,569 |  | ns |
| 16 vs. 1 | -1,046 | -4,739 to 2,648 |  | ns |
| 16 vs. 7 | -0,2265 | -3,920 to 3,467 |  | ns |
| 23 vs. 1 | -0,9214 | -4,615 to 2,772 |  | ns |
| 23 vs. 7 | -0,1023 | -3,795 to 3,591 |  | ns |
| 1 vs. 7 | 0,8191 | -2,874 to 4,512 |  | Ns |
| *ThNAD-MEα* |  |  |  |  |
| 9 vs. 16 | -0,5292 | -0,9437 to -0,1148 | <0.01 | ** |
| 9 vs. 23 | -0,2204 | -0,6349 to 0,1941 |  | ns |
| 9 vs. 1 | -0,3033 | -0,7178 to 0,1111 |  | ns |
| 9 vs. 7 | -0,1978 | -0,6123 to 0,2167 |  | ns |
| 16 vs. 23 | 0,3089 | -0,06185 to 0,6796 |  | ns |
| 16 vs. 1 | 0,2259 | -0,1448 to 0,5966 |  | ns |
| 16 vs. 7 | 0,3314 | -0,03928 to 0,7021 |  | ns |
| 23 vs. 1 | -0,08295 | -0,4537 to 0,2878 |  | ns |
| 23 vs. 7 | 0,02258 | -0,3481 to 0,3933 |  | ns |
| 1 vs. 7 | 0,1055 | -0,2652 to 0,4762 |  | ns |
| *ThNAD-MEβ1* |  |  |  |  |
| 9 vs. 16 | -0,008602 | -0,3793 to 0,3621 |  | ns |
| 9 vs. 23 | 0,002147 | -0,3686 to 0,3729 |  | ns |
| 9 vs. 1 | -0,007697 | -0,3784 to 0,3630 |  | ns |

|  |  |  |  |
| --- | --- | --- | --- |
| 9 vs. 7 | 0,005132 | -0,3656 to 0,3758 | ns |
| 16 vs. 23 | 0,01075 | -0,3600 to 0,3815 | ns |
| 16 vs. 1 | 0,0009052 | -0,3698 to 0,3716 | ns |
| 16 vs. 7 | 0,01373 | -0,3570 to 0,3844 | ns |
| 23 vs. 1 | -0,009844 | -0,3806 to 0,3609 | ns |
| 23 vs. 7 | 0,002986 | -0,3677 to 0,3737 | ns |
| 1 vs. 7 | 0,01283 | -0,3579 to 0,3835 | ns |
| *ThNAD-MEβ2* |  |  |  |
| 9 vs. 16 | -0,03982 | -0,4105 to 0,3309 | ns |
| 9 vs. 23 | -0,01839 | -0,3891 to 0,3523 | ns |
| 9 vs. 1 | -0,02241 | -0,3931 to 0,3483 | ns |
| 9 vs. 7 | 0,005249 | -0,3655 to 0,3760 | ns |
| 16 vs. 23 | 0,02143 | -0,3493 to 0,3921 | ns |
| 16 vs. 1 | 0,01741 | -0,3533 to 0,3881 | ns |
| 16 vs. 7 | 0,04507 | -0,3256 to 0,4158 | ns |
| 23 vs. 1 | -0,004018 | -0,3747 to 0,3667 | ns |
| 23 vs. 7 | 0,02364 | -0,3471 to 0,3943 | ns |
| 1 vs. 7 | 0,02765 | -0,3431 to 0,3984 | ns |

**Supplemental Table 2.** Identification of NAD(P)-ME subunits by MS. (A) Identification of NAD-ME subunits of *G. gynandra* in protein bands of migration levels 1 and 4 in leaf (L) and root (R). (B) Identification of NAD-ME subunits of *C. angustifolia* in protein bands of migration levels 1 and 4 in leaf (L) (C) Identification of NAD-ME subunits of *T. hassleriana* in the protein bands of migration levels 3 in leaf (L) and root (R). (D) Identification of NADP-ME in *G. gynandra*, *C. angustifolia*, and *T. hassleriana* in the protein bands of migration levels 2. The migration level refers to that in Figure 2A. The numbers given refer to the number of peptide spectrum matches. *Number in brackets indicates the identification of two peptides exclusive for ThNAD-MEβ2 in that sample.

**A**
| Migration level | GgNAD-ME |  |  |  |  |  |
| --- | --- | --- | --- | --- | --- | --- |
| | $\alpha$ | | β1 | | β2 | |
|  | L | R | L | R | L | R |
| 1 | 246 | 15 | 19 | 0 | 14 | 24 |
|  | 70 | 18 | 63 | 0 | 7 | 22 |
| 4 | 23 | - | 6 | - | 0 | - |
|  | 29 | - | 13 | - | 0 | - |

**B**
| Migration level | CaNAD-ME |  |  |
| --- | --- | --- | --- |
| | $\alpha$ | β1 | β2 |
|  | L | L | L |
| 1 | 2 | 5 | 3 |
| 4 | 8 | 4 | 0 |

**C**
| Migration level | ThNAD-ME |  |  |  |  |  |
| --- | --- | --- | --- | --- | --- | --- |
| | $\alpha$ | | β1 | | β1/2 | |
|  | L | R | L | R | L | R |
| 3 | 5 | 11 | 9 | 2 | 20 | 14(2*) |
|  | 6 | 17 | 2 | 5 | 7 | 15 |

**D**
| Migration level | NADP-ME |  |  |  |  |  |
| --- | --- | --- | --- | --- | --- | --- |
|  | Gg |  | Ca |  | Th |  |
|  | L | R | L | R | L | R |
| 2 | 17 | 73 | 7 | 11 | 48 | 402 |
|  | 40 | 26 | - | - | 71 | 25 |

**Supplemental Table 3.**
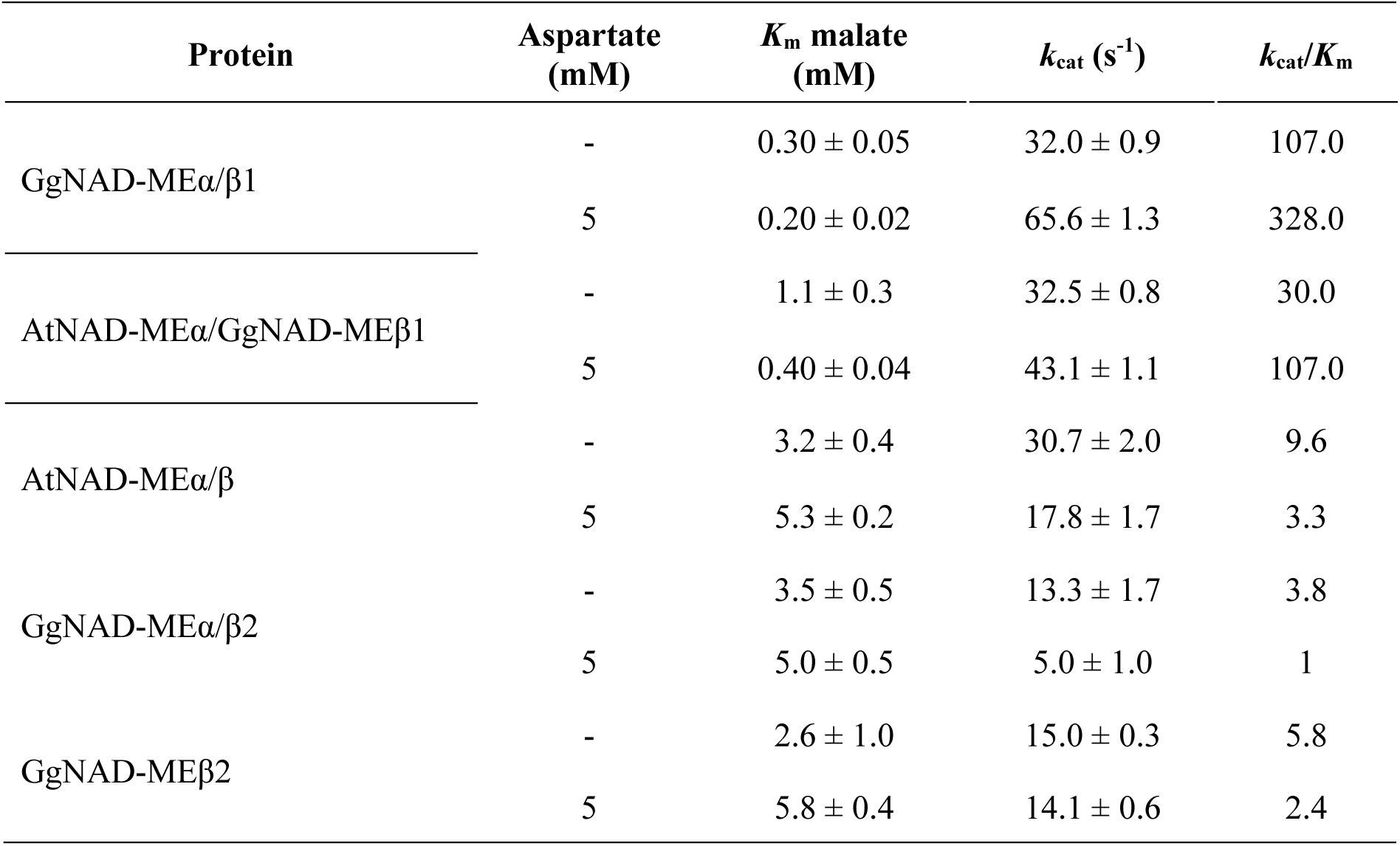
Kinetic parameters of recombinant GgNAD-ME entities in the absence and the presence of aspartate. As GgNAD-MEβ1 is not active, *k*_cat_ was calculated assuming the formation of a heterodimer with only one active site for GgNAD-MEα/β1 and AtNAD-MEα/GgNAD-MEβ1. Three independent enzyme preparations were used in each experiment, each of which was measured in triplicate.

**Supplemental Table 4.**
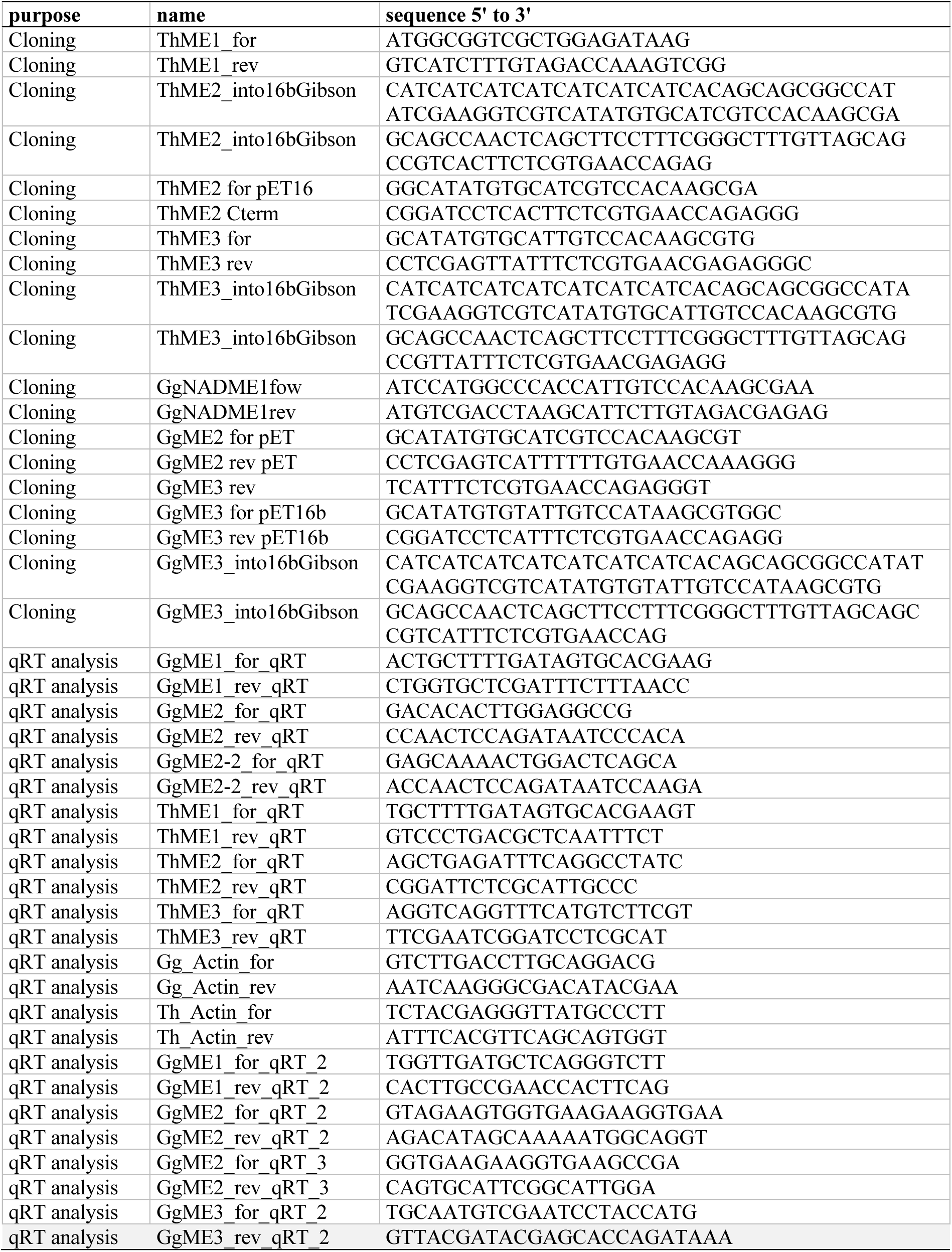
List of primers used in this study.

## SUPPLEMENTAL INFORMATION

### METHODS

#### Induction of protein expression and purification of recombinant NAD-MEs

For induction of protein expression, 400 mL liquid cell cultures containing 100 µg/mL ampicillin (P1) or 100 µg/mL ampicillin and 20 µg/mL gentamicin (P2) were inoculated with freshly grown over-night culture, grown at 37° C for 2.5 h to an OD_600_ of 0.5 (P1) or at 37° C for 3 h to an OD_600_ of 0.8 (P2). Protein expression was induced with 1 mM IPTG. Cells were harvested after 20 h growth at 37° C (P1) or 3 days grown at 12° C (P2) by centrifugation at 4° C and 6000 g, and stored at −20° C until use for a maximum of three month. Purification of His-tagged single proteins was performed using the immobilized metal ion affinity chromatography principle. *E. coli* cells were resuspended in freshly prepared ice-cold lysis buffer (0.2 M NaCl, 20 mM Tris-HCl pH 8.0, 5 mM imidazole, 2 mM phenylmethylsulfonyl fluoride (PMSF), 1 spatula tip lysozyme), incubated for 10 min on ice and sonified for 4 min in 30 s intervals. After centrifugation (15 min, 15000 g, 4° C), supernatant was loaded on a 900 µL nickel-nitrilotriacetic acid (Ni-NTA) agarose (Qiagen, Hilden, Germany) column. Three washing steps with incrementing imidazole concentration were performed (0.2 M NaCl, 20 mM Tris-HCl pH 8.0, and 5, 40, 80 mM (P1) or 5, 40, 60 mM (P2) imidazole) at 4°C with 10 column volumes each until purity of the respective protein was achieved. Proteins were eluted in 2 mL elution buffer (0.2 M NaCl, 20 mM Tris-HCl pH 8.0, 0.5 M imidazole). The elution of NAD-MEα, - β1 and -β2 proteins was verified by Coomassie-stained SDS-PAGE. The expected molecular masses were approximately 67.5, 66.3 and 66.4 kDa for NAD-MEα, -β1 and -β2, respectively.

For purification of co-expressed proteins, cells were resuspended in 80 ml buffer A (20 mM Tris-HCl, pH 7.9, 5 mM imidazole, and 2 mM phenylmethylsulfonyl fluoride), sonicated, and centrifuged for 10 min at 7000 *g* at 4° C. The supernatant was loaded onto a Ni-NTA agarose (Qiagen, Hilden, Germany) column previously equilibrated with buffer A. The co-expressed proteins were eluted with buffer A containing 200 mM imidazole. The co-elution of both NAD-MEα and -β proteins was verified by Coomassie-stained SDS-PAGE. The expected molecular masses were 75-82 kDa for the pET32-expressed proteins (58-65 kDa corresponding to the NAD-ME + 17 kDa the His-Trx•Tag) and 58-65 kDa in the case of NAD-MEs expressed in the pET29 vector. The equimolar ratio of the co-purified proteins was estimated by densitometric analysis of the bands obtained after SDS-PAGE of the eluted fractions.

The co-purified fusion NAD-MEs were incubated with thrombin protease (1:100) for 2 h at 15° C to remove the N-terminus encoded by the pET32 vector. The proteins were further purified by using gel-chromatography Sephadex G-50 column equilibrated with buffer B (50 mM MES pH 6.5, 5 mM MnCl_2_, 5 mM dithiothreitol and, and 20 % (v/v) glycerol). Purified enzymes were stored at −80° C in bluffer B with 50 % glycerol as was previously described (Tronconi et al., 2008).

#### Mass spectrometry

*G. gynandra* and *T. hassleriana* samples were analyzed on a QExactive Plus mass spectrometer. Briefly, full scans were carried out at a resolution of 70000 (scan range 200 to 2000 m/z, profile mode, automatic gain control target value 3E6, maximum injection time 50 ms). Subsequently, up to 20 2-5fold precursors were selected (4 m/z isolation window), fragmented by higher-energy collisional dissociation (HCD) and analyzed at a resolution of 17500 (available scan range 200 to 2000 m/z, centroid mode, automatic gain control target value 1E5, maximum injection time 50 ms), the dynamic exclusion was 10 s. Parameter for the *G. gynandra* leaf band migration level 1 sample 2 were as indicated above with the following modifications: MS1 scan range 350 – 2000 m/z, MS1 maximum injection time 80 ms; isolation window 2 m/z; MS2 maximum injection time 60 ms, top 10 method, 100 s dynamic exclusion.

The *C. angustifolia* proteins were analyzed on a Lumos Fusion hybrid instrument (Thermo Fisher Scientific) with the following parameters: MS1 (Orbitrap) resolution 120000, MS1 scan range 200-2000 m/z, MS1 automatic gain control target 400000, MS1 maximum injection time 60 ms, 1.6 m/z isolation window (charge states 2-7 included), HCD fragmentation, MS2 (linear ion trap) scan rate rapid, MS2 automatic gain control target 400000, MS2 maximum injection time 150 ms, cycle time 2 s, 60 s dynamic exclusion.

Peptide and protein searches were carried out within the Proteome Discoverer 2.4.1.15 framework (Thermo Fisher Scientific). MS Amanda 2.0 was used as search engine followed by a fixed value peptide spectrum matches (PSM) validator node and peptides and proteins accepted at a false discovery rate of 1%. PSM were reported only for unambiguously assignable peptides and if a minum of two PSM were identified for one protein. For MS Amanda, following settings were applied: trypic cleavage specificity, maximum of 2 missed cleavages, carbamidomethylation at cysteines as fixed and protein N-terminal acetylation and methionine oxidation as variable modifications, MS1 tolerance 5 ppm, MS2 tolerance 0.02 Da (0.4 Da for *C. angustifolia* proteins). Acquired spectra were matched against transcriptome derived *G. gynandra* (32832 entries) and *T. hassleriana* (28718 entries) protein databases based on the transcriptome datasets of Külahoglu et al. (2014). For identification of *C. angustifolia* proteins, a custom database was created using the *G. gynandra* transcriptome and adding the known NAD-ME coding sequences.

#### Generation of comparative models

We used SWISS-MODEL (Waterhouse et al., 2018) to build comparative models of the α, β1, and β2 isoforms of GgNAD-ME as well as the α and β_1_ isoform of ThNAD-ME. All models were built using PDB ID 1PJ3 (Tao et al., 2003) as a template structure (sequence identities: 42.0-42.8 %, coverage: > 88 %). After superimposing the protein structures, the coordinates of NAD^+^, L-malate, and Mg^2+^ in PDB ID 1PJ2 (Tao et al., 2003) were copied to the comparative models. Crystal waters within 4.0 Å of any of the placed molecules were taken from PDB ID 1PJ2, too. Molecules bound to the template outside the active site were not considered. Using the protein preparation wizard of MAESTRO (Schrödinger Release 2018-1: Maestro; Schrödinger, LLC, New York, NY, 2018), we capped the termini and protonated the structures according to pH 7.0 using Epik (Schrödinger Release 2018-1: Epik; Schrödinger, LLC, New York, NY, 2018). Unusual protonation states (protonated His or protonated Asp and Glu) were visually inspected; the protons were afterward removed. The lysin residue interacting with the 2-OH group of L-malate was set to a deprotonated, uncharged state to simulate the state of the enzyme before the oxidation of L-malate. All hydrogens were minimized using the OPLS3e force field while keeping all heavy atoms fixed. Afterward, crystal water molecules from PDB ID 1PJ3 further away than 2.2 Å away from L-malate, NAD^+^, and Mg^2+^ were placed in the structures. The protonated structures of the monomeric isoforms were used to build the GgNAD-MEα/α homodimer as well as the GgNAD-MEα/β1, GgNAD-MEα/β2, and ThNAD-MEα/β1 heterodimers by superimposing the monomers onto 1PJ3.

#### Preparation of molecular dynamics simulations

tLEaP from the Amber20 software package (AMBER 2020; University of California, San Francisco, 2020) was used to prepare the complexes for the molecular dynamics (MD) simulations. The complexes were solvated in an octahedral box of TIP3P water (Jorgensen et al., 1983), leaving at least 11.0 Å to any edge of the box. Random water molecules were replaced by sodium to neutralize the charges. ff14SB (Maier et al., 2015) was used for the proteins. For L-malate and NAD^+^, GAFF2 parameters were used, and atomic point charges were derived using the RESP procedure (Bayly et al., 1993) from structures optimized with Gaussian 16 (Gaussian 16 Rev. A.03; Gaussian, Inc., Wallingford CT, 2016) at the HF/6-31G* level of theory. The Mg^2+^ cations in the catalytic site were modeled employing a cationic dummy atom model (Jiang et al., 2015).

#### Thermalization of molecular dynamics simulations

Initially, we minimized the structures three times through 3000 steps of steepest descent followed by 2000 steps using the conjugate gradient algorithm. First, C_α_ atoms, the metal center, and the heavy atoms of the substrates were held fixed with a force constant of 5 kcal mol^−1^ Å^−2^. Second, only water molecules and C_α_ atoms were held fixed and, third, no restraints were used. Keeping the positions of the C_α_ atoms, the metal center, and the heavy atoms of the substrates fixed with a force constant of 2 kcal mol^−1^ Å^−2^, the systems were heated to 300 K over 25 ps of NVT-MD (constant number of particles, volume, and temperature) and simulated for further 25 ps. For density adaptation, 450 ps of NPT-MD (constant number of particles, pressure, and temperature) at 1.0 bar were performed. Afterward, the restraints on the atomic positions were gradually removed over 1,500 ns of NPT-MD. Finally, the systems were simulated for a further 1,000 ps without any restraints.

#### Quantifying changes of biomolecular stability along “constraint dilution trajectories”

The changes of the biomolecular stability along the “constraint dilution trajectories” are quantified in neighbor stability maps on a residue-wise level, yielding a potential energy for each rigid contact (*rc*) formed between pairs of residues (Rathi et al., 2015; Wifling et al., 2019). A residue’s chemical potential energy due to non-covalent bonding is obtained as the sum of all short-range rigid contacts the residue is involved in (Pfleger et al., 2017; Pfleger et al., 2013) (Eq. S1).

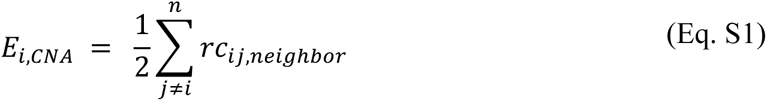

The CNA software was applied on conformational ensembles of 2500 snapshots extracted from the MD simulations of each system. Prior to the rigidity analysis, water molecules, sodium ions, and cap groups were removed from the structures, while NAD^+^, L-malate, and the Mg^2+^ ion were kept. Visualization of the molecular structures and the representation of the rigid contacts was done with PyMOL (DeLano, 2002).

#### Assessing the impact of individual substitutions in the GgNAD-MEβ1 isoform on the biomolecular rigidity and flexibility

To assess the impact of individual substitutions in the GgNAD-MEβ1 isoform on the biomolecular rigidity and flexibility, we applied a perturbation approach implemented in CNA (Pfleger et al., 2017). Ground state network topologies were generated from conformational ensembles as described above. Then, all covalent and non-covalent restraints associated with the sidechain of a selected residue are removed, yielding the perturbed state, where the selected residue is stripped down to an alanine residue. As both states are generated from the same conformational ensemble, any changes in the local stability can be related directly to the *in silico* alanine substitution (Pfleger et al., 2017). Decomposing the free energy associated with the change in biomolecular stability on a per-residue level (Δ*G_i_*_,CNA_) allows monitoring how the perturbation effect percolates through the enzyme structure, indicating the structural coupling of remote areas to the substitution site. By performing the same perturbation on corresponding residues in GgNAD-MEβ1 and GgNAD-MEβ2, ΔΔ*G_i_*_,CNA_ is obtained for each residue *i*, which shows the changes in rigidity for the substitution between the isoforms (Eq. S2).

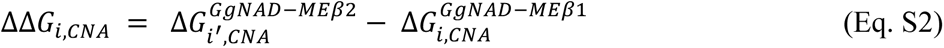

Here, *i* and *i’* are corresponding residues in GgNAD-MEβ1 and GgNAD-MEβ2 according to a sequence alignment (Tronconi et al., 2020).

